# Super-resolved spatial transcriptomics by deep data fusion

**DOI:** 10.1101/2020.02.28.963413

**Authors:** Ludvig Bergenstråhle, Bryan He, Joseph Bergenstråhle, Alma Andersson, Joakim Lundeberg, James Zou, Jonas Maaskola

## Abstract

In situ RNA capturing has made it possible to record histology and spatial gene expression from the same tissue section. Here, we introduce a method that combines data from both modalities to infer super-resolved full-transcriptome expression maps. Our method unravels transcriptional heterogeneity in micrometer-scale anatomical features and enables image-based in silico spatial transcriptomics without hybridization or sequencing.

---

Spatial transcriptomics allows researchers to study cell behavior in the spatial domain and has been used to describe cellular organization in the hippocampus [1], to characterize intra-tumor heterogeneity in human breast [2], pancreatic [3], and prostate cancer [4], to analyze spatial dynamics during embryonic cardiogenesis [5], and in many other contexts.

Experimental methods for spatial transcriptomics fall on a spectrum that trades resolution and molecular sensitivity for multiplexing capacity. On one end of the spectrum, methods based on in situ sequencing [6, 7] or hybridization [8, 9, 10] typically have high resolution and high sensitivity but are difficult to multiplex over many genes, limiting their usefulness in exploring transcriptome-wide interactions. On the other end, methods based on in situ RNA capturing (ISC) using poly(dT) probes [2, 11, 12] target all poly-adenylated transcripts simultaneously but have lower resolution and sensitivity, limiting their usefulness in studying detailed expression patterns.

To overcome the limitations of current spatial transcriptomics methods, we propose a deep generative model of spatial expression data. Our model casts spatial gene expression and histological image data as observable effects of a latent tissue state (Fig. 1a, Methods). By fusing low-sensitivity, low-resolution ISC expression data with high-resolution histological image data, we infer denoised full-transcriptome spatial gene expression at the same resolution as the image data. Additionally, our model can be applied to samples that lack expression data, making it possible to predict spatial gene expression without hybridization or sequencing.

**Figure 1:**
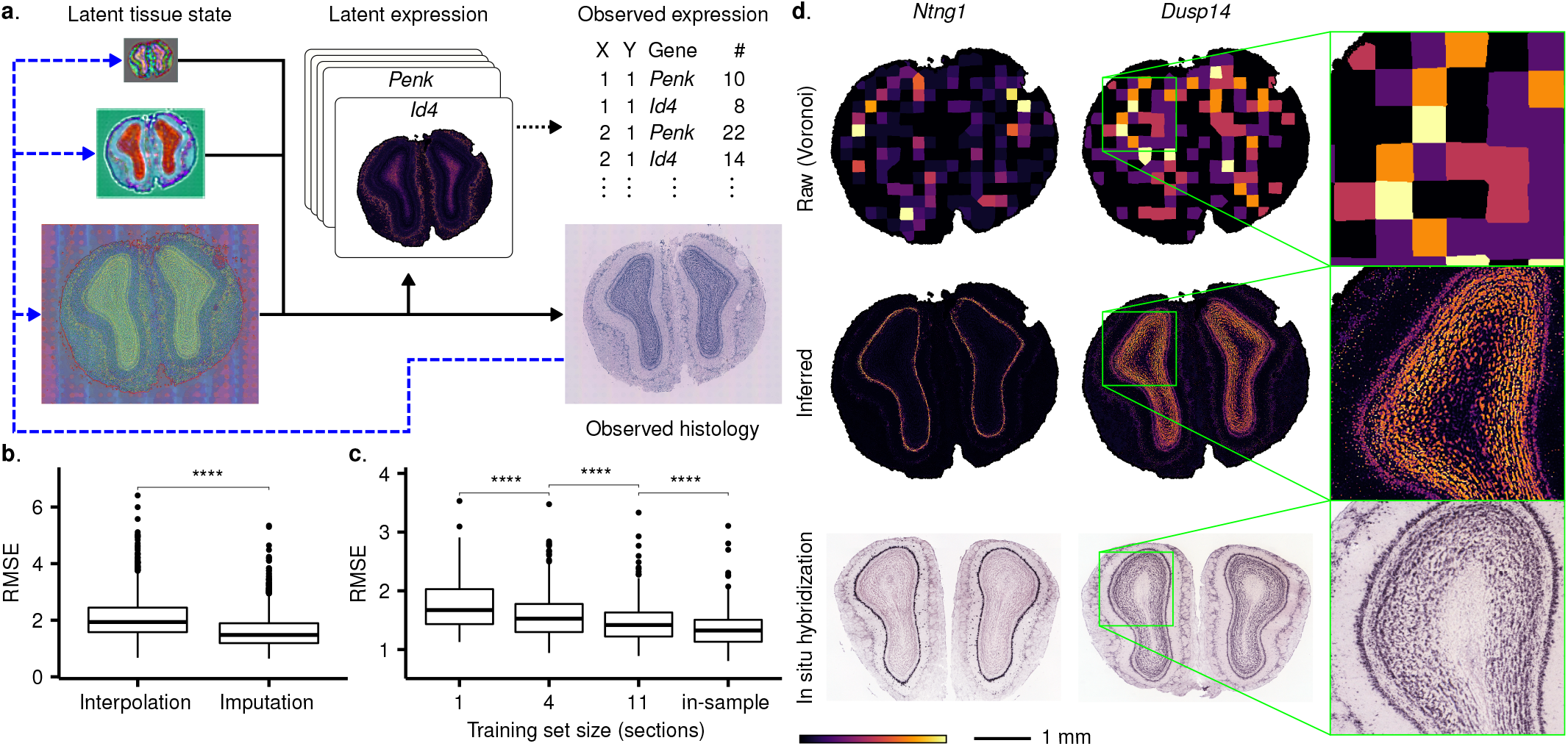
Conceptual overview and performance evaluation. (a) Histological image and expression data are modeled as effects of a latent spatial tissue state. The tissue state has multiple resolutions, capturing both global and local anatomical features, and is mapped through a generator network (black solid arrows) to histological image and high-resolution, latent expression data. The latent expression data is linked to the observed expression data by summation (black dotted arrow). Inference is amortized using a recognition network (blue dashed arrows) that maps the observed image data to the latent tissue state. (b), (c) Root-mean-square error (RMSE) of (b) imputation compared to pixel-wise, zero-order interpolation and (c) out-of-sample prediction over different training set sizes. Count values are normalized to mean one in each measurement location. Asterisks (****) indicate significance at the *p* ≤ 0.0001 level using a two-sided Wilcoxon signed-rank test. (d) Comparison of inferred high-resolution expression maps to in situ hybridization reference data from the Allen Mouse Brain Atlas.

We model the latent tissue state over multiple spatial resolutions, capturing both global and local anatomical features. Inference of the latent state and corresponding high-resolution expression data is based on ideas from the literature on variational autoencoders [13, 14]. Importantly, while optimizing model parameters, we jointly learn a recognition neural network that maps the image data to the variational parameters of the latent state. As a result, the inferred posterior of the latent state is not kept in memory but recomputed for each mini-batch, allowing our method to scale to arbitrarily large datasets.

To evaluate the performance of the proposed method, we study a dataset [2] consisting of 12 sections from the mouse olfactory bulb. First, we test in-sample performance by dropping 50% of all measurement locations and imputing the missing expression data. We compare the results to a pixel-wise interpolation scheme that fills in missing data with the expression of the closest non-missing location and find that our method achieves a 24 % lower median root-mean-square error (Fig. 1b).

We next test out-of-sample performance by holding out an entire section from the training set and predicting its expression data from the image data. We find that ground truth expression patterns are faithfully reproduced (Fig. S1) and that accuracy approaches in-sample performance as more sections are included in the training set (Fig. 1c), demonstrating that our method can be applied to histological images that do not have associated ISC data.

Finally, we compare inferred expression to in situ hybridization data from the Allen Mouse Brain Atlas [15]. Overall, inferred expression closely matches the reference data (Fig. 1d and Fig. S2). For example, expression of Ntng1 in the mitral cell layer (MCL) and of Dusp14 in the MCL and granule layer are accurately replicated (Fig. 1d). In contrast, the raw data is too coarse to resolve the same expression patterns.

We use our method to study detailed anatomical structures in the mouse olfactory bulb and in human breast cancer. Both datasets display fine-grained expression heterogeneity (Figs. 2a and 2b), which can be quantified in terms of differential expression (Methods).

**Figure 2:**
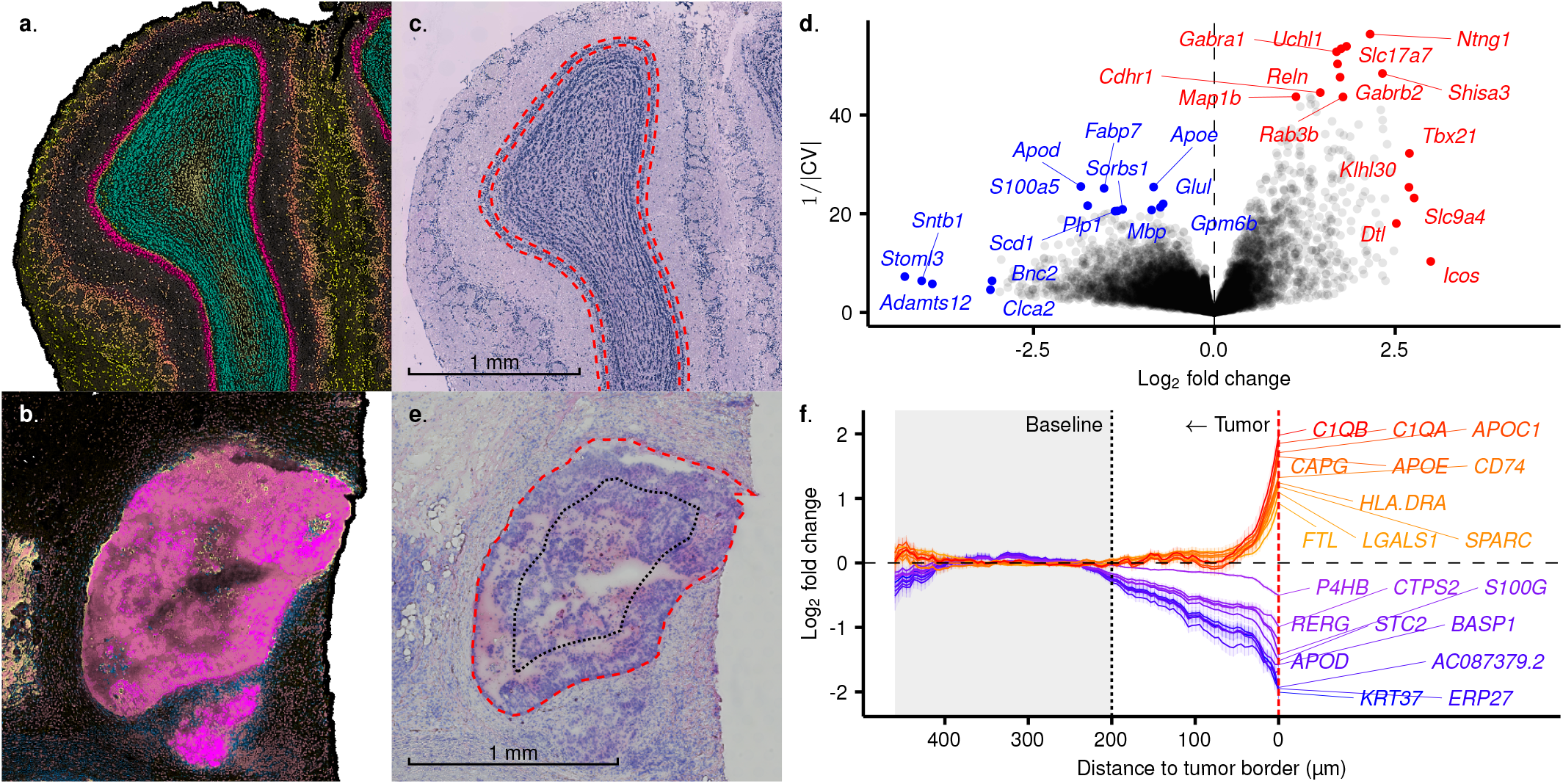
Characterization of transcriptional heterogeneity in detailed anatomical structures. (a), (b) Summarized latent gene expression in (a) the mouse olfactory bulb and (b) a human ductal carcinoma in situ (DCIS) lesion. Colors indicate anatomical areas with distinct transcriptional phenotypes according to the inferred tissue state (Methods). (c) Annotation of the mitral cell layer (MCL) profiled in (d). (d) Differential expression in the MCL compared to the other layers of the mouse olfactory bulb. (e) Annotation of the DCIS lesion profiled in (f). Red dashed line: Tumor border. Black dotted line: Baseline boundary, 200 μm from the tumor border. (f) Differential expression compared to baseline as a function of proximity to the tumor border for the 10 most up- and downregulated genes at distance zero. Lines show posterior means and ribbons indicate uncertainty (±2 standard deviations).

First, we profile the MCL of the olfactory bulb (Fig. 2c) and find several strongly up- and downregulated genes (Fig. 2d). To verify our results, we sort the genes by the inverted coefficient of variation of their posterior log_2_ fold change and find that 40 out of the 100 most upregulated genes are among 229 markers for the MCL identified in a recent single-cell RNA-sequencing study [16] (one-sided hypergeometric test *p*-value: 1.66 × 10^-47^). Meanwhile, the 20–50 μm thickness of the MCL prohibits it from being isolated in the raw data, which measures expression over areas with a diameter of 100 μm. We conclude that the proposed method successfully deconvolves mixed expression signals by integrating expression patterns across anatomical areas that share morphological features.

Next, we study spatial dynamics in a ductal carcinoma in situ (DCIS) lesion from the breast cancer dataset by profiling transcriptome gradients between the inner area of the tumor and its outermost edge (Fig. 2e). We find several genes related to immune activity and tumor progression to be upregulated at the border of the tumor (Fig. 2f). For example, the complement component 1q, composed of the C1QA, C1QB, and C1QC subcomponents, have been shown to promote angiogenesis and tumor growth in the tumor microenvironment [17]. Similarly, CD74 is a known marker for metastatic tumor growth in breast cancer [18] and is being investigated as a potential target for antibody-drug conjugate therapies in blood cancers. The proximity of CD74 expression to the tumor edge could have important implications for the accessibility of CD74 expressing cells in similar therapies for DCIS. However, further studies are needed to validate this finding.

Consistent with the above results, the pathways that are enriched for the 100 most upregulated genes at the tumor border include, for example, extracellular structure organization (*p*-value: 2.10 × 10^-18^), immune system processes (*p*-value: 1.37 × 10^-11^), blood vessel development (*p*-value: 8.51 × 10^-6^), and cell migration regulation (*p*-value: 8.98 × 10^-6^) (Table S1).

Critically, while the distance between measurement locations in the raw expression data is 100 μm, several differentially expressed genes become upregulated first within 50 μm of the tumor border (Fig. 2f). We conclude that it is only by learning a high-resolution state of the underlying transcriptional anatomy of the tissue that it becomes possible to fully resolve the detailed expression landscape describing these genes. Determining its precise topology is paramount to understanding cellular interactions at the microscale and in developing effective treatments for a wide range of diseases.

In summary, we have presented a deep generative model for spatial data fusion. We combine ISC expression data with histological image data to infer super-resolved full-transcriptome spatial gene expression. The proposed method exposes spatial contingencies that are difficult to discern in the raw expression data and can characterize differential expression in micrometer-scale anatomical features. Moreover, our method can predict spatial gene expression from histology in samples that lack ISC data, thereby providing a means for image-based in *silico spatial transcriptomics* (ISST).

We envision future work to enable ISST on a grander scale. Given a sufficiently large and diverse training set, it may be possible to learn universal models that can predict spatial gene expression without hybridization or sequencing in any tissue. ISST could make very large spatial transcriptomics projects economically viable, unlock spatial gene expression in vast databases of histology images, or be used to verify the integrity of data from experimental methods.

## Acknowledgments

This work was made possible by generous support from the Knut and Alice Wallenberg foundation, the Erling-Persson family foundation, the Swedish Cancer Society, the Swedish Foundation for Strategic Research, and the Swedish Research Council.

## Author contributions

L.B. and J.M. designed the method. L.B. implemented the method and wrote the paper. B.H., J.B., and A.A. provided valuable feedback and contributed to the analysis. J.M., J.Z., and J.L. supervised the project.

## Methods

### Statistical model

We model the spatial expression data, *X_n_*, and histological image data, *I_n_*, of each sample *n* as effects of an underlying spatial tissue state, *Z_n_*. We assume the conditional distribution of the image data *I* to be Gaussian, and, following previous work [1] on RNA bulk sequencing data, we assume the conditional distribution of the expression data *X* to be negative binomial. The rate of the latter is factorized into *M* metagenes, parameterized by a gene loading matrix *L*. The parameters of the conditional distributions are mapped from the latent tissue state *Z* through a convolutional generator network *G* with learnable parameters *θ*.

Formally, for all samples *n*, pixel coordinates (*x, y*), genes *g*, and image channels *c*, we model the data generating process as follows:

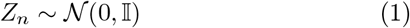

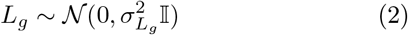

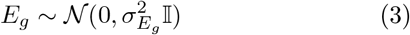

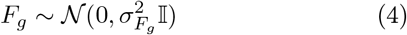

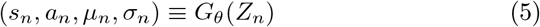

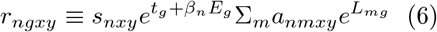

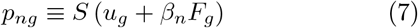

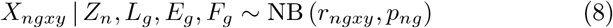

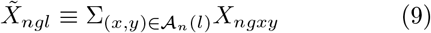

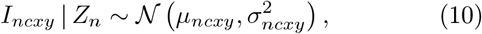

where *S* is the logistic function, *β_n_* is a row vector of indicator variables specifying group membership, and 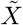 is the observed expression data at location *l* covering the area 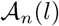. The fixed effects *E* and *F* can be used to control for batch effects or to characterize differential expression between sample groups.

During inference, we collapse the model by integrating out the latent expression *X*, which replaces Eqs. (8) and (9) with

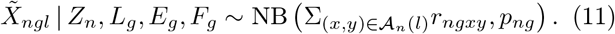

### Inference

We use variational inference to approximate the posterior of the latent variables 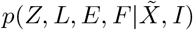 with a tractable distribution *q_ϕ_*(*Z, L, E, F*). The variational parameters *φ* and the parameters *θ* of the generator network are found by minimizing the Kullback-Leibler divergence from *q_φ_* to the posterior, which is equivalent to maximizing the evidence lower bound (ELBO),

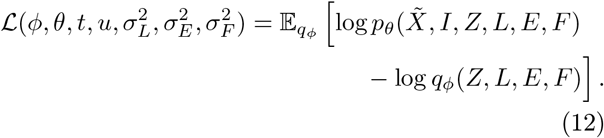

We use a mean-field diagonal Gaussian variational distribution

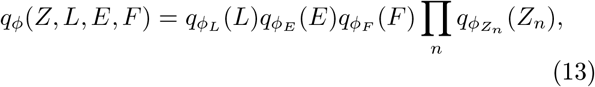

where the parameters *φ_Z_n__* are encoded by a convolutional recognition network *R* with weights *φ_Z_* applied to the image data: *φ_Z_n__* ≡ *R_φ__Z_* (In).

We update the parameters 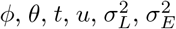, and 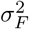 by gradient ascent on the objective (12) using the Adam optimizer [2]. Following [3], gradient estimates are obtained by reparameterizing the latent variables as a function of auxiliary parameter-free noise. Briefly, letting

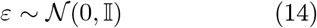

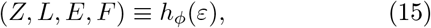

where *h_φ_* is an appropriate shift-and-scale transformation, we can reformulate Eq. (12) as an expectation with respect to *ε* by relying on the law of the unconscious statistician. This makes it straightforward to rewrite the gradient of Eq. (12) as an expectation,

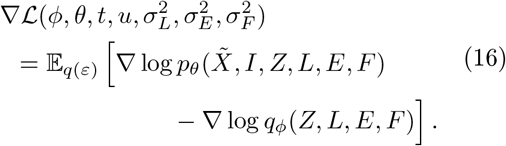

We approximate (16) using a single Monte Carlo sample for each update step and train on patches extracted from the dataset.

The dataset is augmented with random rotations, scaling, and shearing. The image data is further augmented with random color jitter.

### Architecture

To efficiently capture both global and local anatomical features, we model the latent tissue state *Z* over multiple resolutions. The recognition and generator networks *G* and *R* together form an architecture similar to U-Net [4] with the variational distribution of the latent state for each resolution inserted at the corresponding skip connection (Fig. S3).

### Model selection

To select the number of metagenes *M* in the model, we implement a drop-and-split strategy that runs in parallel to inference. Briefly, we start out with *M* = 1 metagenes. At fixed intervals, we estimate the ELBO (12) with and without each of the *M* metagenes. Metagenes that contribute to the ELBO are split into two new metagenes that inherit parameters from their parent while non-contributing metagenes are dropped.

### High-resolution gene expression maps

We infer denoised latent gene expression by estimating the posterior distribution of

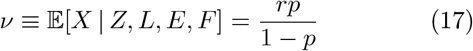

with *N* Monte Carlo samples drawn from the variational distribution:

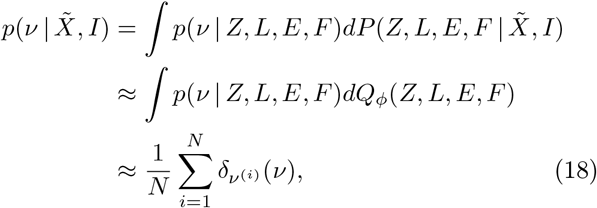

where *P* and *Q_φ_* denote the cumulative distribution functions of the corresponding lower-case densities and *δ*_*ν*(*i*)_ is the Dirac delta function centered at the i:th sample of ν.

We compute gene expression maps as the mean of the point-mass mixture (18),

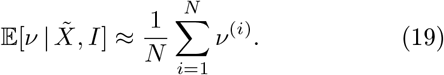

To predict latent gene expression in an unseen sample *n*′, we approximate

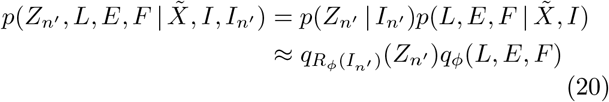

and estimate 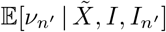 similar to Eq. (19).

### Differential expression analysis

We consider the log_2_ conditional mean expression of an area 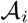,

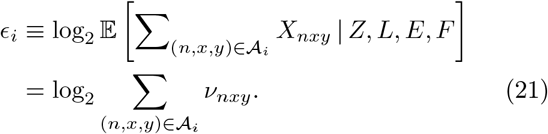

The posterior distribution of the normalized log_2_ fold change of a gene *g* between the areas 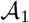 and 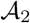,

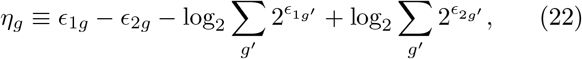

is estimated analogous to Eq. (18). Mean and variance estimates are computed on the resultant point-mass mixture:

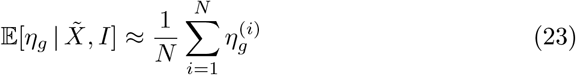

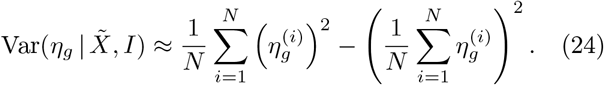

### Summarized expression maps

To visualize transcriptional anatomy, we estimate the posterior mean metagene activity a and pixel-wise scale s similar to Eq. (19). We project a onto its first three principal components and append -s along the channel axis. We then apply a channel-wise affine transformation to map all values into [0, 1]. The resulting coordinates are used as CMYK-encoded color values.

### Pathway analysis

Pathway analyses are conducted using g:Profiler [5] with the GO:BP database [6, 7]. Reported *p*-values are adjusted with the g:SCS procedure provided by g:Profiler.

### Relationship to prior work

Our work extends previous research on spatial models of transcriptomics data. Notably, SpatialDE [8] and SPARK [9] model spatial transcriptomics data using Gaussian processes to detect spatially variable genes. However, neither method makes use of histological information or can be used to infer high-resolution expression data. NovoSpaRc [10] reconstructs the spatial organization of single cells by solving an optimal transport problem. While novoSpaRc can identify zonated genes from single-cell data, accurate inference of spatial expression patterns requires information about the spatial configuration of marker genes. Several other methods [11, 12, 13, 14] exist for fusing single cell with in situ sequencing or hybridization data.

The contribution of our work is threefold: First, we have shown that histological image data is highly informative of spatial expression patterns in tissues. Second, we provide an integrative model of in situ capturing spatial transcriptomics. Our model fuses spatial gene expression data with high-resolution image data, thereby making it possible to study full-transcriptome expression heterogeneity in detailed anatomical structures. Third, we have demonstrated the feasibility of predicting expression in un- squenced samples using only their histological image data. We believe image-based in silico spatial transcriptomics to be a promising future research topic.

## Data availability

The mouse olfactory bulb dataset was obtained from the spatial research group’s website: https://www.spatialresearch.org. The breast cancer dataset was obtained from 10X Genomics: https://support.10xgenomics.com/spatial-gene-expression/datasets/.

## Code availability

We have implemented the proposed method in the Pyro probabilistic programming language [15]. The code is available under the MIT license at https://github.com/ludvb/xfuse.

