## Supplementary materials for "Super-resolved spatial transcriptomics by deep data fusion"

2020-03-12

### Supplementary Figures

S1 In silico spatial transcriptomics . . . 2
S2 Comparison to in situ hybridization reference data . . . 3
S3 Architecture . . . 4

### Supplementary Tables

S1 Upregulated pathways . . . 5

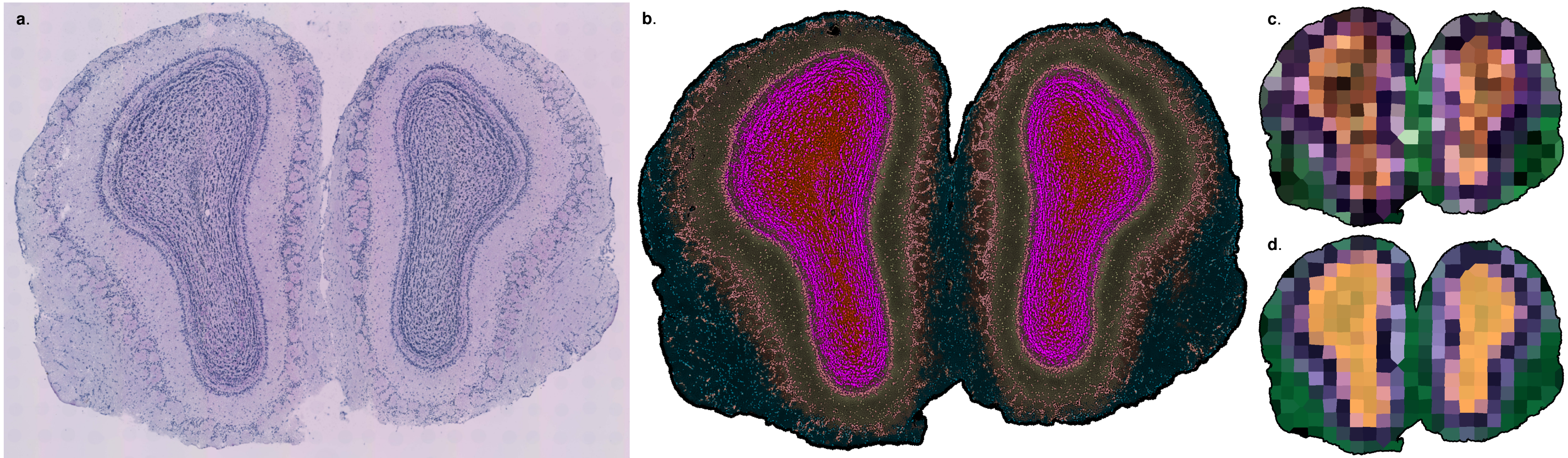


**Figure S1**: In silico spatial transcriptomics. (a) Histological image data of the held-out section. (b) Summarized expression map of the predicted metagene expression associated with (a). (c), (d) Comparison of summarized expression maps constructed from (c) normalized log ground truth gene expression in the held-out section and (d) normalized log predicted gene expression at the ground truth measurement locations using data from (b).

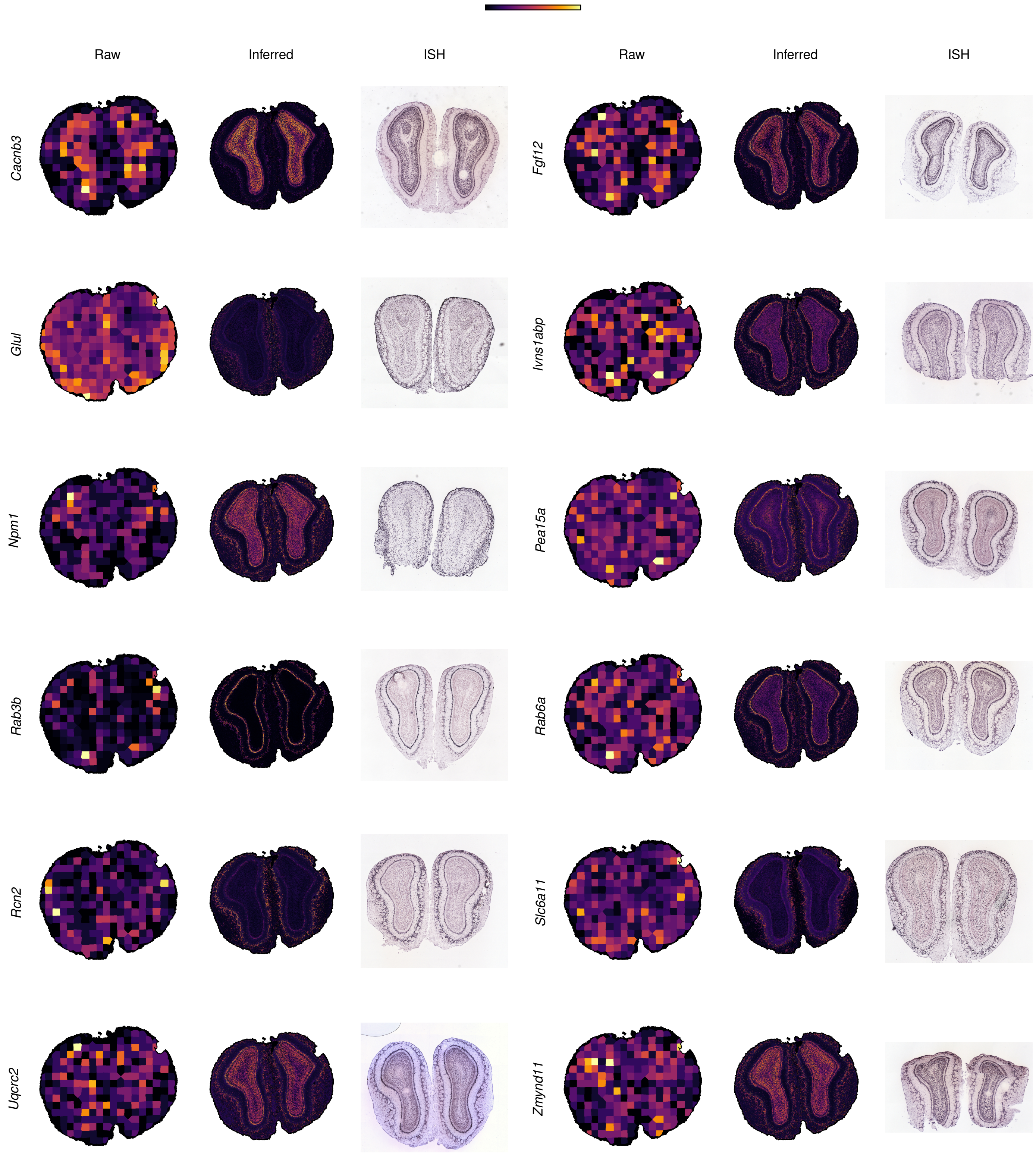


**Figure S2**: Comparison of inferred high-resolution expression maps to in situ hybridization reference data from the Allen Mouse Brain Atlas. Random samples from the 1000 most expressed genes. Raw: Raw expression data (Voronoi tesselation). Inferred: Inferred high-resolution expression maps. ISH: In situ hybridization reference data.

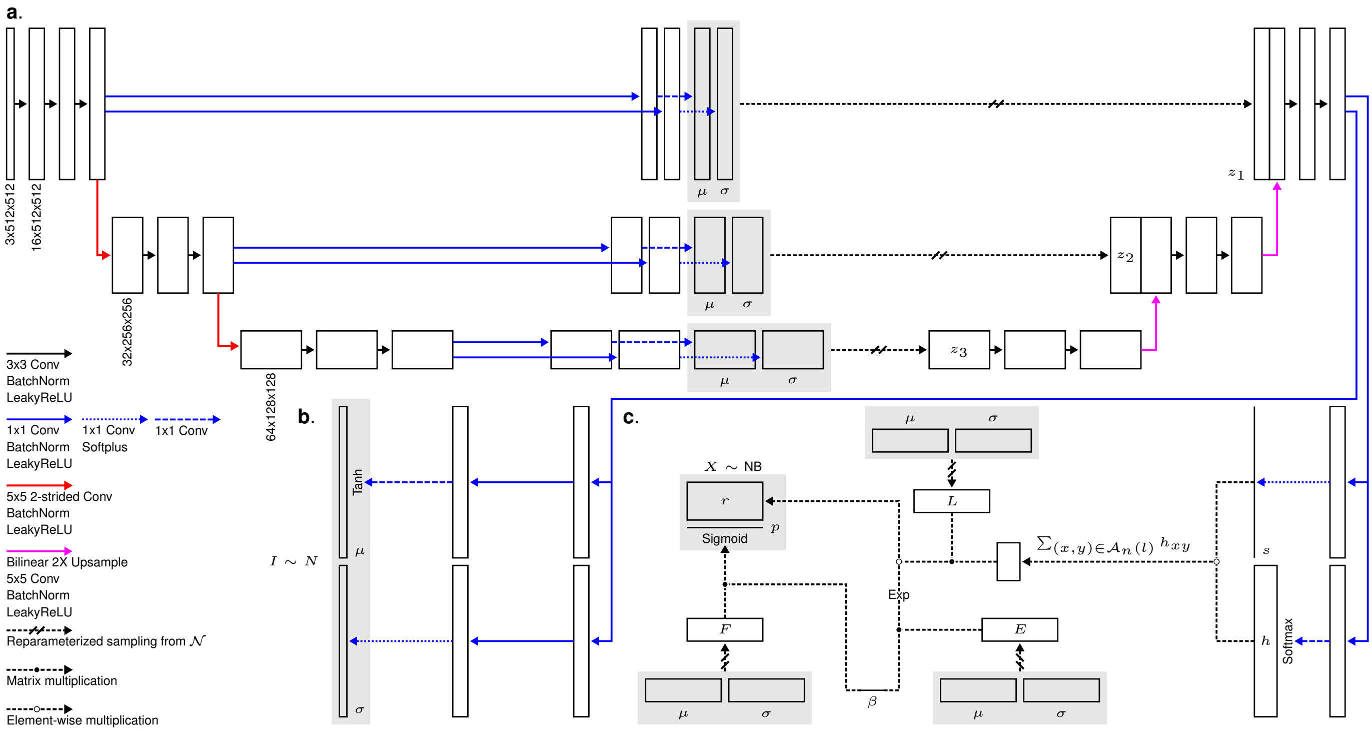


**Figure S3**: Architecture. (a) Fusion network. (b) Image data decoder. (c) Expression data decoder. Volume dimensions and number of down- and upsampling steps are exemplative.

**Table S1**: Upregulated pathways at DCIS tumor border

| Biological process | Adjusted *p*-value | Intersecting genes |
|---|---|---|
| Extracellular Structure Organization | $2.10 \times 10^{-18}$ | *APOE*, *APOC1*, *SPARC*, *COL6A1*, *SFRP2*, *AEBP1*, *COL1A1*, *LUM*, *A2M*, *POSTN*, *ITGB2*, *BGN*, *COL1A2*, *FN1*, *COL3A1*, *COL6A2*, *TIMP1*, *MMP11*, *CTSS*, *EMILIN1*, *DCN*, *COMP*, *LRP1*, *COL6A3*, *HSPG2*, *FBN1* |
| Extracellular Matrix Organization | $2.81 \times 10^{-17}$ | *SPARC*, *COL6A1*, *SFRP2*, *AEBP1*, *COL1A1*, *LUM*, *A2M*, *POSTN*, *ITGB2*, *BGN*, *COL1A2*, *FN1*, *COL3A1*, *COL6A2*, *TIMP1*, *MMP11*, *CTSS*, *EMILIN1*, *DCN*, *COMP*, *LRP1*, *COL6A3*, *HSPG2*, *FBN1* |
| Immune System Process | $1.37 \times 10^{-11}$ | *APOE*, *CD74*, *C1QB*, *FTL*, *C1QA*, *LGALS1*, *C1QC*, *CRIP1*, *CYBA*, *SFRP2*, *CRIP2*, *COL1A1*, *MDK*, *A2M*, *GPNMB*, *PTMS*, *LYZ*, *MUC1*, *CTSD*, *VIM*, *ITGB2*, *C3*, *COL1A2*, *FN1*, *CTSB*, *TYROBP*, *PSAP*, *PYCARD*, *FAU*, *COL3A1*, *SERPING1*, *CORO1A*, *CTSS*, *LGMN*, *EMILIN1*, *FLNA*, *B2M*, *CD68*, *CD14*, *FCER1G*, *C1S*, *LRP1*, *RARRES2*, *ACTB*, *TAPBP*, *SPI1*, *PSMB8*, *FBN1* |
| Defense Response | $2.15 \times 10^{-11}$ | *APOE*, *CD74*, *C1QB*, *C1QA*, *C1QC*, *CYBA*, *MDK*, *A2M*, *LYZ*, *MUC1*, *VIM*, *ITGB2*, *C3*, *FN1*, *CTSB*, *TYROBP*, *PYCARD*, *FAU*, *SERPING1*, *CORO1A*, *TIMP1*, *CTSS*, *LGMN*, *FLNA*, *B2M*, *CD68*, *CD14*, *SERPINF1*, *FCER1G*, *C1S*, *LRP1*, *RARRES2*, *HSPG2*, *LSP1*, *PSMB8*, *TGM2* |
| Cell Activation | $1.66 \times 10^{-10}$ | *APOE*, *CD74*, *FTL*, *C1QA*, *LGALS1*, *CYBA*, *COL1A1*, *MDK*, *GPNMB*, *LYZ*, *CTSD*, *ITGB2*, *C3*, *COL1A2*, *CTSB*, *TYROBP*, *PSAP*, *PYCARD*, *COL3A1*, *CORO1A*, *TIMP1*, *CTSS*, *EMILIN1*, *FLNA*, *B2M*, *COMP*, *CD68*, *CD14*, *FCER1G*, *LRP1*, *ACTB*, *SPI1* |
| Regulation Of Immune System Process | $1.93 \times 10^{-10}$ | *APOE*, *CD74*, *C1QB*, *C1QA*, *LGALS1*, *C1QC*, *CYBA*, *COL1A1*, *MDK*, *A2M*, *GPNMB*, *MUC1*, *ITGB2*, *C3*, *COL1A2*, *CTSB*, *TYROBP*, *PYCARD*, *COL3A1*, *SERPING1*, *CORO1A*, *CTSS*, *LGMN*, *EMILIN1*, *B2M*, *CD68*, *CD14*, *FCER1G*, *C1S*, *RARRES2*, *ACTB*, *SPI1*, *PSMB8*, *FBN1* |
| Immune Response | $3.76 \times 10^{-10}$ | *APOE*, *CD74*, *C1QB*, *FTL*, *C1QA*, *LGALS1*, *C1QC*, *CRIP1*, *CYBA*, *COL1A1*, *MDK*, *A2M*, *LYZ*, *MUC1*, *CTSD*, *VIM*, *ITGB2*, *C3*, *COL1A2*, *CTSB*, *TYROBP*, *PSAP*, *PYCARD*, *FAU*, *COL3A1*, *SERPING1*, *CORO1A*, *CTSS*, *LGMN*, *B2M*, *CD68*, *CD14*, *FCER1G*, *C1S*, *LRP1*, *RARRES2*, *ACTB*, *TAPBP*, *PSMB8* |
| Regulated Exocytosis | $6.07 \times 10^{-10}$ | *FTL*, *SPARC*, *CYBA*, *A2M*, *LYZ*, *CTSD*, *ITGB2*, *ISLR*, *C3*, *FN1*, *CTSB*, *TYROBP*, *PSAP*, *PYCARD*, *SERPING1*, *CORO1A*, *TIMP1*, *CTSS*, *FLNA*, *B2M*, *CD68*, *CD14*, *FCER1G*, *RARRES2* |
| Response To External Stimulus | $5.88 \times 10^{-9}$ | *APOE*, *CD74*, *C1QB*, *C1QA*, *SPARC*, *C1QC*, *CYBA*, *SFRP2*, *COL1A1*, *MDK*, *A2M*, *GPNMB*, *LYZ*, *POSTN*, *MUC1*, *VIM*, *ITGB2*, *C3*, *CTSB*, *TYROBP*, *PYCARD*, *FAU*, *SEMA3F*, *COL3A1*, *TYMP*, *SERPING1*, *CORO1A*, *CTSS*, *LGMN*, *FLNA*, *GFRA1*, *DCN*, *B2M*, *CD68*, *CD14*, *SERPINF1*, *FCER1G*, *C1S*, *LRP1*, *RARRES2*, *PSMB8*, *TGM2* |
| Collagen Fibril Organization | $9.42 \times 10^{-9}$ | *SFRP2*, *AEBP1*, *COL1A1*, *LUM*, *COL1A2*, *COL3A1*, *MMP11*, *EMILIN1*, *COMP* |
| Immune Effector Process | $9.99 \times 10^{-9}$ | *CD74*, *C1QB*, *FTL*, *C1QA*, *LGALS1*, *C1QC*, *CYBA*, *MDK*, *A2M*, *LYZ*, *CTSD*, *ITGB2*, *C3*, *CTSB*, *TYROBP*, *PSAP*, *PYCARD*, *SERPING1*, *CORO1A*, *CTSS*, *FLNA*, *B2M*, *CD68*, *CD14*, *FCER1G*, *C1S*, *LRP1*, *ACTB* |
| Exocytosis | $1.07 \times 10^{-8}$ | *FTL*, *SPARC*, *CYBA*, *A2M*, *LYZ*, *CTSD*, *ITGB2*, *ISLR*, *C3*, *FN1*, *CTSB*, *TYROBP*, *PSAP*, *PYCARD*, *SERPING1*, *CORO1A*, *TIMP1*, *CTSS*, *FLNA*, *B2M*, *CD68*, *CD14*, *FCER1G*, *RARRES2* |
| Response To Stress | $3.34 \times 10^{-8}$ | *APOE*, *CD74*, *C1QB*, *C1QA*, *LGALS1*, *SPARC*, *C1QC*, *CRIP1*, *CYBA*, *SFRP2*, *COL1A1*, *MDK*, *A2M*, *LYZ*, *POSTN*, *MUC1*, *VIM*, *ITGB2*, *C3*, *COL1A2*, *FN1*, *CTSB*, *TYROBP*, *PSAP*, *PYCARD*, *FAU*, *COL3A1*, *SERPING1*, *CORO1A*, *TIMP1*, *CTSS*, *LGMN*, *FLNA*, *DCN*, *B2M*, *COMP*, *CD68*, *CD14*, *SERPINF1*, *FCER1G*, *C1S*, *LRP1*, *RARRES2*, *ACTB*, *ELOB*, *HSPG2*, *LSP1*, *PSMB8*, *TGM2* |
| Secretion By Cell | $3.35 \times 10^{-8}$ | *APOE*, *FTL*, *SPARC*, *CYBA*, *MDK*, *A2M*, *LYZ*, *POSTN*, *CTSD*, *ITGB2*, *ISLR*, *C3*, *FN1*, *CTSB*, *TYROBP*, *PSAP*, *PYCARD*, *SELENOM*, *SERPING1*, *CORO1A*, *TIMP1*, *CTSS*, *FLNA*, *B2M*, *COMP*, *CD68*, *CD14*, *FCER1G*, *LRP1*, *RARRES2* |
| Secretion | $5.14 \times 10^{-8}$ | *APOE*, *CD74*, *FTL*, *SPARC*, *CYBA*, *MDK*, *A2M*, *LYZ*, *POSTN*, *CTSD*, *ITGB2*, *ISLR*, *C3*, *FN1*, *CTSB*, *TYROBP*, *PSAP*, *PYCARD*, *SELENOM*, *SERPING1*, *CORO1A*, *TIMP1*, *CTSS*, *FLNA*, *B2M*, *COMP*, *CD68*, *CD14*, *FCER1G*, *LRP1*, *RARRES2* |
| Export From Cell | $6.96 \times 10^{-8}$ | *APOE*, *FTL*, *SPARC*, *CYBA*, *MDK*, *A2M*, *LYZ*, *POSTN*, *CTSD*, *ITGB2*, *ISLR*, *C3*, *FN1*, *CTSB*, *TYROBP*, *PSAP*, *PYCARD*, *SELENOM*, *SERPING1*, *CORO1A*, *TIMP1*, *CTSS*, *FLNA*, *B2M*, *COMP*, *CD68*, *CD14*, *FCER1G*, *LRP1*, *RARRES2* |
| Innate Immune Response | $8.42 \times 10^{-8}$ | *APOE*, *C1QB*, *C1QA*, *C1QC*, *CYBA*, *A2M*, *MUC1*, *VIM*, *ITGB2*, *C3*, *CTSB*, *TYROBP*, *PYCARD*, *FAU*, *SERPING1*, *CORO1A*, *CTSS*, *LGMN*, *B2M*, *CD14*, *FCER1G*, *C1S*, *RARRES2*, *PSMB8* |

| Biological process | Adjusted *p*-value | Intersecting genes |
|---|---|---|
| Defense Response To Other Organism | $9.40 \times 10^{-8}$ | *APOE*, *C1QB*, *C1QA*, *C1QC*, *CYBA*, *A2M*, *LYZ*, *MUC1*, *VIM*, *ITGB2*, *C3*, *CTSB*, *TYROBP*, *PYCARD*, *FAU*, *SERPING1*, *CORO1A*, *CTSS*, *LGMN*, *FLNA*, *B2M*, *CD14*, *FCER1G*, *C1S*, *RARRES2*, *PSMB8* |
| Regulation Of Immune Response | $1.23 \times 10^{-7}$ | *APOE*, *CD74*, *C1QB*, *C1QA*, *C1QC*, *CYBA*, *COL1A1*, *A2M*, *MUC1*, *ITGB2*, *C3*, *COL1A2*, *CTSB*, *TYROBP*, *PYCARD*, *COL3A1*, *SERPING1*, *CTSS*, *LGMN*, *B2M*, *CD14*, *FCER1G*, *C1S*, *ACTB*, *PSMB8* |
| Response To Other Organism | $1.53 \times 10^{-7}$ | *APOE*, *C1QB*, *C1QA*, *SPARC*, *C1QC*, *CYBA*, *A2M*, *LYZ*, *MUC1*, *VIM*, *ITGB2*, *C3*, *CTSB*, *TYROBP*, *PYCARD*, *FAU*, *SERPING1*, *CORO1A*, *CTSS*, *LGMN*, *FLNA*, *DCN*, *B2M*, *CD68*, *CD14*, *FCER1G*, *C1S*, *RARRES2*, *PSMB8* |
| Response To External Biotic Stimulus | $1.58 \times 10^{-7}$ | *APOE*, *C1QB*, *C1QA*, *SPARC*, *C1QC*, *CYBA*, *A2M*, *LYZ*, *MUC1*, *VIM*, *ITGB2*, *C3*, *CTSB*, *TYROBP*, *PYCARD*, *FAU*, *SERPING1*, *CORO1A*, *CTSS*, *LGMN*, *FLNA*, *DCN*, *B2M*, *CD68*, *CD14*, *FCER1G*, *C1S*, *RARRES2*, *PSMB8* |
| Negative Regulation Of Multicellular Organismal Process | $1.81 \times 10^{-7}$ | *APOE*, *CD74*, *APOC1*, *LGALS1*, *SPARC*, *C1QC*, *CYBA*, *SFRP2*, *MDK*, *GPNMB*, *VIM*, *FN1*, *TYROBP*, *PYCARD*, *SEMA3F*, *COL3A1*, *SERPING1*, *TIMP1*, *LGMN*, *EMILIN1*, *DCN*, *B2M*, *SERPINF1*, *INPP5J*, *LRP1*, *HSPG2*, *FBN1* |
| Cellular Response To Chemical Stimulus | $2.19 \times 10^{-7}$ | *APOE*, *CD74*, *LGALS1*, *SPARC*, *CRIP1*, *COL6A1*, *LAPTM5*, *CYBA*, *CPNE7*, *SFRP2*, *COL1A1*, *MDK*, *POSTN*, *MUC1*, *VIM*, *ITGB2*, *COL1A2*, *FN1*, *CTSB*, *PSAP*, *PYCARD*, *COL3A1*, *CORO1A*, *TIMP1*, *CTSS*, *LGMN*, *EMILIN1*, *FLNA*, *DCN*, *B2M*, *COMP*, *CD68*, *CD14*, *SERPINF1*, *FCER1G*, *LRP1*, *RARRES2*, *ACTB*, *ELOB*, *VSTM2A*, *SPI1*, *PSMB8*, *FBN1* |
| Response To Biotic Stimulus | $2.25 \times 10^{-7}$ | *APOE*, *C1QB*, *C1QA*, *SPARC*, *C1QC*, *CYBA*, *A2M*, *LYZ*, *MUC1*, *VIM*, *ITGB2*, *C3*, *CTSB*, *TYROBP*, *PYCARD*, *FAU*, *SERPING1*, *CORO1A*, *CTSS*, *LGMN*, *FLNA*, *DCN*, *B2M*, *CD68*, *CD14*, *FCER1G*, *C1S*, *RARRES2*, *PSMB8* |
| Leukocyte Mediated Immunity | $2.66 \times 10^{-7}$ | *CD74*, *C1QB*, *FTL*, *C1QA*, *C1QC*, *CYBA*, *LYZ*, *CTSD*, *ITGB2*, *C3*, *CTSB*, *TYROBP*, *PSAP*, *PYCARD*, *SERPING1*, *CORO1A*, *CTSS*, *B2M*, *CD68*, *CD14*, *FCER1G*, *C1S* |
| Response To Organic Substance | $2.78 \times 10^{-7}$ | *APOE*, *CD74*, *LGALS1*, *SPARC*, *CRIP1*, *COL6A1*, *LAPTM5*, *CYBA*, *SFRP2*, *COL1A1*, *LUM*, *MDK*, *POSTN*, *MUC1*, *VIM*, *ITGB2*, *COL1A2*, *FN1*, *CTSB*, *PSAP*, *PYCARD*, *COL3A1*, *COL6A2*, *CORO1A*, *TIMP1*, *CTSS*, *LGMN*, *EMILIN1*, *FLNA*, *DCN*, *B2M*, *COMP*, *CD68*, *CD14*, *SERPINF1*, *FCER1G*, *LRP1*, *RARRES2*, *ACTB*, *VSTM2A*, *SPI1*, *PSMB8*, *FBN1* |
| Positive Regulation Of Immune System Process | $4.99 \times 10^{-7}$ | *CD74*, *C1QB*, *C1QA*, *LGALS1*, *C1QC*, *CYBA*, *MDK*, *A2M*, *MUC1*, *ITGB2*, *C3*, *CTSB*, *TYROBP*, *PYCARD*, *SERPING1*, *CORO1A*, *CTSS*, *LGMN*, *B2M*, *CD14*, *FCER1G*, *C1S*, *RARRES2*, *ACTB*, *PSMB8* |
| Animal Organ Development | $7.01 \times 10^{-7}$ | *CD74*, *C1QB*, *LGALS1*, *SPARC*, *C1QC*, *CRIP1*, *COL6A1*, *SFRP2*, *CRIP2*, *COL1A1*, *LUM*, *MDK*, *GPNMB*, *VIM*, *SLCO2B1*, *BGN*, *CTHRC1*, *COL1A2*, *FN1*, *CTSB*, *TYROBP*, *PSAP*, *SELENOM*, *SEMA3F*, *COL3A1*, *COL6A2*, *TIMP1*, *EMILIN1*, *FLNA*, *DCN*, *B2M*, *COMP*, *SERPINF1*, *FCER1G*, *RPL10*, *LRP1*, *RARRES2*, *ACTB*, *COL6A3*, *HSPG2*, *SPI1*, *PSMB8*, *TGM2*, *FBN1* |
| Cellular Response To Organic Substance | $8.83 \times 10^{-7}$ | *CD74*, *LGALS1*, *SPARC*, *COL6A1*, *LAPTM5*, *CYBA*, *SFRP2*, *COL1A1*, *POSTN*, *MUC1*, *VIM*, *ITGB2*, *COL1A2*, *FN1*, *CTSB*, *PSAP*, *PYCARD*, *COL3A1*, *CORO1A*, *TIMP1*, *CTSS*, *LGMN*, *EMILIN1*, *FLNA*, *DCN*, *B2M*, *COMP*, *CD68*, *CD14*, *SERPINF1*, *FCER1G*, *LRP1*, *RARRES2*, *ACTB*, *VSTM2A*, *SPI1*, *PSMB8*, *FBN1* |
| System Development | $9.09 \times 10^{-7}$ | *APOE*, *CD74*, *C1QB*, *C1QA*, *LGALS1*, *SPARC*, *C1QC*, *CRIP1*, *COL6A1*, *SFRP2*, *CRIP2*, *COL1A1*, *LUM*, *MDK*, *GPNMB*, *POSTN*, *VIM*, *SLCO2B1*, *ITGB2*, *C3*, *BGN*, *CTHRC1*, *COL1A2*, *FN1*, *CTSB*, *TYROBP*, *PSAP*, *SELENOM*, *SEMA3F*, *COL3A1*, *COL6A2*, *TYMP*, *TIMP1*, *EMILIN1*, *FLNA*, *GFRA1*, *DCN*, *B2M*, *COMP*, *SERPINF1*, *INPP5J*, *FCER1G*, *RPL10*, *LRP1*, *RARRES2*, *ACTB*, *COL6A3*, *HSPG2*, *SPI1*, *PSMB8*, *TGM2*, *FBN1* |
| Cellular Component Organization | $9.97 \times 10^{-7}$ | *APOE*, *CD74*, *APOC1*, *C1QB*, *C1QA*, *CAPG*, *LGALS1*, *SPARC*, *C1QC*, *COL6A1*, *CYBA*, *SFRP2*, *AEBP1*, *COL1A1*, *LUM*, *MDK*, *A2M*, *REPS2*, *POSTN*, *MUC1*, *VIM*, *ITGB2*, *C3*, *BGN*, *CTHRC1*, *COL1A2*, *FN1*, *TYROBP*, *PSAP*, *PYCARD*, *SEMA3F*, *COL3A1*, *COL6A2*, *TYMP*, *CORO1A*, *TIMP1*, *MMP11*, *CTSS*, *LGMN*, *EMILIN1*, *FLNA*, *GFRA1*, *DCN*, *B2M*, *COMP*, *CD14*, *SERPINF1*, *INPP5J*, *FCER1G*, *RPL10*, *LRP1*, *ACTB*, *TAPBP*, *ELOB*, *COL6A3*, *HSPG2*, *SPI1*, *SPARCL1*, *TGM2*, *FBN1* |
| Platelet Degranulation | $1.13 \times 10^{-6}$ | *SPARC*, *A2M*, *ISLR*, *FN1*, *PSAP*, *SERPING1*, *TIMP1*, *FLNA*, *FCER1G*, *RARRES2* |
| Response To Chemical | $1.31 \times 10^{-6}$ | *APOE*, *CD74*, *C1QA*, *LGALS1*, *SPARC*, *CRIP1*, *COL6A1*, *LAPTM5*, *CYBA*, *CPNE7*, *SFRP2*, *COL1A1*, *LUM*, *MDK*, *GPNMB*, *POSTN*, *MUC1*, *VIM*, *ITGB2*, *COL1A2*, *FN1*, *CTSB*, *PSAP*, *PYCARD*, *SELENOM*, *SEMA3F*, *COL3A1*, *COL6A2*, *TYMP*, *CORO1A*, *TIMP1*, *CTSS*, *LGMN*, *EMILIN1*, *FLNA*, *GFRA1*, *DCN*, *B2M*, *COMP*, *CD68*, *CD14*, *SERPINF1*, *FCER1G*, *LRP1*, *RARRES2*, *ACTB*, *ELOB*, *VSTM2A*, *SPI1*, *PSMB8*, *FBN1* |
| Response To Wounding | $1.46 \times 10^{-6}$ | *APOE*, *LGALS1*, *SPARC*, *COL1A1*, *MDK*, *A2M*, *POSTN*, *COL1A2*, *FN1*, *TYROBP*, *COL3A1*, *SERPING1*, *TIMP1*, *FLNA*, *DCN*, *COMP*, *FCER1G*, *LRP1*, *ACTB* |

| Biological process | Adjusted *p*-value | Intersecting genes |
|---|---|---|
| Regulation Of Developmental Process | $2.02 \times 10^{-6}$ | *APOE*, *CD74*, *LGALS1*, *SPARC*, *C1QC*, *SFRP2*, *COL1A1*, *MDK*, *GPNMB*, *POSTN*, *VIM*, *ITGB2*, *C3*, *CTHRC1*, *FN1*, *TYROBP*, *SEMA3F*, *COL3A1*, *TYMP*, *CORO1A*, *TIMP1*, *MMP11*, *LGMN*, *EMILIN1*, *FLNA*, *DCN*, *B2M*, *COMP*, *SERPINF1*, *INPP5J*, *LRP1*, *RARRES2*, *VSTM2A*, *HSPG2*, *SPI1*, *PSMB8*, *FBN1* |
| Cell Activation Involved In Immune Response | $2.04 \times 10^{-6}$ | *FTL*, *LGALS1*, *CYBA*, *MDK*, *LYZ*, *CTSD*, *ITGB2*, *C3*, *CTSB*, *TYROBP*, *PSAP*, *PYCARD*, *CORO1A*, *CTSS*, *B2M*, *CD68*, *CD14*, *FCER1G*, *LRP1* |
| Cellular Component Organization Or Biogenesis | $3.50 \times 10^{-6}$ | *APOE*, *CD74*, *APOC1*, *C1QB*, *C1QA*, *CAPG*, *LGALS1*, *SPARC*, *C1QC*, *COL6A1*, *CYBA*, *SFRP2*, *AEBP1*, *COL1A1*, *LUM*, *MDK*, *A2M*, *REPS2*, *POSTN*, *MUC1*, *VIM*, *ITGB2*, *C3*, *BGN*, *CTHRC1*, *COL1A2*, *FN1*, *TYROBP*, *PSAP*, *PYCARD*, *SEMA3F*, *COL3A1*, *COL6A2*, *TYMP*, *CORO1A*, *TIMP1*, *MMP11*, *CTSS*, *LGMN*, *EMILIN1*, *FLNA*, *GFRA1*, *DCN*, *B2M*, *COMP*, *CD14*, *SERPINF1*, *INPP5J*, *FCER1G*, *RPL10*, *LRP1*, *ACTB*, *TAPBP*, *ELOB*, *COL6A3*, *HSPG2*, *SPI1*, *SPARCL1*, *TGM2*, *FBN1* |
| Developmental Process | $3.73 \times 10^{-6}$ | *APOE*, *CD74*, *C1QB*, *C1QA*, *LGALS1*, *SPARC*, *C1QC*, *CRIP1*, *COL6A1*, *SFRP2*, *CRIP2*, *COL1A1*, *LUM*, *MDK*, *A2M*, *GPNMB*, *POSTN*, *VIM*, *SLCO2B1*, *ITGB2*, *C3*, *BGN*, *CTHRC1*, *COL1A2*, *FN1*, *CTSB*, *TYROBP*, *PSAP*, *SELENOM*, *SEMA3F*, *COL3A1*, *COL6A2*, *TYMP*, *SERPING1*, *CORO1A*, *TIMP1*, *MMP11*, *LGMN*, *EMILIN1*, *FLNA*, *GFRA1*, *DCN*, *B2M*, *COMP*, *CD68*, *SERPINF1*, *INPP5J*, *FCER1G*, *RPL10*, *LRP1*, *RARRES2*, *ACTB*, *COL6A3*, *VSTM2A*, *HSPG2*, *SPI1*, *PSMB8*, *TGM2*, *FBN1* |
| Myeloid Leukocyte Activation | $4.14 \times 10^{-6}$ | *CD74*, *FTL*, *C1QA*, *CYBA*, *LYZ*, *CTSD*, *ITGB2*, *C3*, *CTSB*, *TYROBP*, *PSAP*, *PYCARD*, *CTSS*, *B2M*, *CD68*, *CD14*, *FCER1G*, *SPI1* |
| Inflammatory Response | $7.65 \times 10^{-6}$ | *APOE*, *C1QA*, *CYBA*, *MDK*, *LYZ*, *ITGB2*, *C3*, *FN1*, *TYROBP*, *PYCARD*, *TIMP1*, *CD68*, *CD14*, *SERPINF1*, *FCER1G*, *LRP1*, *RARRES2*, *HSPG2*, *TGM2* |
| Regulation Of Multicellular Organismal Process | $8.05 \times 10^{-6}$ | *APOE*, *CD74*, *APOC1*, *LGALS1*, *SPARC*, *C1QC*, *CYBA*, *SFRP2*, *COL1A1*, *LUM*, *MDK*, *GPNMB*, *POSTN*, *VIM*, *ITGB2*, *C3*, *CTHRC1*, *FN1*, *TYROBP*, *PYCARD*, *SEMA3F*, *COL3A1*, *TYMP*, *SERPING1*, *TIMP1*, *LGMN*, *EMILIN1*, *FLNA*, *DCN*, *B2M*, *COMP*, *CD14*, *SERPINF1*, *INPP5J*, *FCER1G*, *LRP1*, *HSPG2*, *SPI1*, *PSMB8*, *FBN1* |
| Blood Vessel Development | $8.51 \times 10^{-6}$ | *APOE*, *SPARC*, *SFRP2*, *COL1A1*, *MDK*, *GPNMB*, *ITGB2*, *C3*, *COL1A2*, *FN1*, *COL3A1*, *TYMP*, *EMILIN1*, *DCN*, *COMP*, *SERPINF1*, *LRP1*, *HSPG2*, *SPI1* |
| Regulation Of Cell Migration | $8.98 \times 10^{-6}$ | *APOE*, *CD74*, *SPARC*, *SFRP2*, *COL1A1*, *MDK*, *GPNMB*, *POSTN*, *FN1*, *PYCARD*, *SEMA3F*, *COL3A1*, *CORO1A*, *TIMP1*, *LGMN*, *EMILIN1*, *FLNA*, *DCN*, *SERPINF1*, *LRP1*, *RARRES2* |
| Negative Regulation Of Developmental Process | $9.20 \times 10^{-6}$ | *APOE*, *CD74*, *LGALS1*, *SPARC*, *C1QC*, *SFRP2*, *MDK*, *POSTN*, *VIM*, *SEMA3F*, *COL3A1*, *TIMP1*, *MMP11*, *LGMN*, *EMILIN1*, *DCN*, *B2M*, *SERPINF1*, *INPP5J*, *LRP1*, *HSPG2*, *FBN1* |
| Leukocyte Degranulation | $1.06 \times 10^{-5}$ | *FTL*, *CYBA*, *LYZ*, *CTSD*, *ITGB2*, *C3*, *CTSB*, *TYROBP*, *PSAP*, *PYCARD*, *CORO1A*, *CTSS*, *B2M*, *CD68*, *CD14*, *FCER1G* |
| Leukocyte Activation Involved In Immune Response | $1.29 \times 10^{-5}$ | *FTL*, *LGALS1*, *CYBA*, *MDK*, *LYZ*, *CTSD*, *ITGB2*, *C3*, *CTSB*, *TYROBP*, *PSAP*, *PYCARD*, *CORO1A*, *CTSS*, *B2M*, *CD68*, *CD14*, *FCER1G* |
| Response To Endogenous Stimulus | $1.29 \times 10^{-5}$ | *APOE*, *SPARC*, *COL6A1*, *CYBA*, *SFRP2*, *COL1A1*, *MDK*, *POSTN*, *VIM*, *ITGB2*, *COL1A2*, *CTSB*, *COL3A1*, *CORO1A*, *TIMP1*, *CTSS*, *LGMN*, *EMILIN1*, *FLNA*, *COMP*, *CD68*, *SERPINF1*, *FCER1G*, *LRP1*, *RARRES2*, *ACTB*, *VSTM2A*, *FBN1* |
| Multicellular Organism Development | $1.61 \times 10^{-5}$ | *APOE*, *CD74*, *C1QB*, *C1QA*, *LGALS1*, *SPARC*, *C1QC*, *CRIP1*, *COL6A1*, *SFRP2*, *CRIP2*, *COL1A1*, *LUM*, *MDK*, *GPNMB*, *POSTN*, *VIM*, *SLCO2B1*, *ITGB2*, *C3*, *BGN*, *CTHRC1*, *COL1A2*, *FN1*, *CTSB*, *TYROBP*, *PSAP*, *SELENOM*, *SEMA3F*, *COL3A1*, *COL6A2*, *TYMP*, *TIMP1*, *MMP11*, *EMILIN1*, *FLNA*, *GFRA1*, *DCN*, *B2M*, *COMP*, *SERPINF1*, *INPP5J*, *FCER1G*, *RPL10*, *LRP1*, *RARRES2*, *ACTB*, *COL6A3*, *HSPG2*, *SPI1*, *PSMB8*, *TGM2*, *FBN1* |
| Positive Regulation Of Immune Response | $1.63 \times 10^{-5}$ | *CD74*, *C1QB*, *C1QA*, *C1QC*, *CYBA*, *A2M*, *MUC1*, *ITGB2*, *C3*, *CTSB*, *PYCARD*, *SERPING1*, *CTSS*, *LGMN*, *B2M*, *CD14*, *FCER1G*, *C1S*, *ACTB*, *PSMB8* |
| Vasculature Development | $1.75 \times 10^{-5}$ | *APOE*, *SPARC*, *SFRP2*, *COL1A1*, *MDK*, *GPNMB*, *ITGB2*, *C3*, *COL1A2*, *FN1*, *COL3A1*, *TYMP*, *EMILIN1*, *DCN*, *COMP*, *SERPINF1*, *LRP1*, *HSPG2*, *SPI1* |
| Cell Adhesion | $1.96 \times 10^{-5}$ | *CD74*, *LGALS1*, *COL6A1*, *SFRP2*, *COL1A1*, *MDK*, *GPNMB*, *POSTN*, *MUC1*, *ITGB2*, *ISLR*, *FN1*, *PYCARD*, *COL3A1*, *COL6A2*, *CORO1A*, *EMILIN1*, *FLNA*, *COMP*, *LRP1*, *ACTB*, *COL6A3*, *SPARCL1*, *TGM2*, *FBN1* |
| Vesicle-Mediated Transport | $2.05 \times 10^{-5}$ | *APOE*, *APOC1*, *FTL*, *SPARC*, *CYBA*, *A2M*, *LYZ*, *CTSD*, *ITGB2*, *ISLR*, *C3*, *FN1*, *CTSB*, *TYROBP*, *PSAP*, *PYCARD*, *SERPING1*, *CORO1A*, *TIMP1*, *CTSS*, *FLNA*, *B2M*, *CD68*, *CD14*, *FCER1G*, *LRP1*, *RARRES2*, *ACTB*, *TAPBP*, *HSPG2*, *TGM2* |
| Activation Of Immune Response | $2.08 \times 10^{-5}$ | *C1QB*, *C1QA*, *C1QC*, *CYBA*, *A2M*, *MUC1*, *ITGB2*, *C3*, *CTSB*, *PYCARD*, *SERPING1*, *CTSS*, *LGMN*, *CD14*, *FCER1G*, *C1S*, *ACTB*, *PSMB8* |
| Cardiovascular System Development | $2.10 \times 10^{-5}$ | *APOE*, *SPARC*, *SFRP2*, *COL1A1*, *MDK*, *GPNMB*, *ITGB2*, *C3*, *COL1A2*, *FN1*, *COL3A1*, *TYMP*, *EMILIN1*, *DCN*, *COMP*, *SERPINF1*, *LRP1*, *HSPG2*, *SPI1* |

| Biological process | Adjusted *p*-value | Intersecting genes |
|---|---|---|
| Connective Tissue Development | $2.11 \times 10^{-5}$ | *CRIP1*, *COL6A1*, *SFRP2*, *COL1A1*, *LUM*, *MDK*, *BGN*, *SELENOM*, *COL6A2*, *TIMP1*, *COMP*, *COL6A3* |
| Biological Adhesion | $2.16 \times 10^{-5}$ | *CD74*, *LGALS1*, *COL6A1*, *SFRP2*, *COL1A1*, *MDK*, *GPNMB*, *POSTN*, *MUC1*, *ITGB2*, *ISLR*, *FN1*, *PYCARD*, *COL3A1*, *COL6A2*, *CORO1A*, *EMILIN1*, *FLNA*, *COMP*, *LRP1*, *ACTB*, *COL6A3*, *SPARCL1*, *TGM2*, *FBN1* |
| Neutrophil Degranulation | $2.33 \times 10^{-5}$ | *FTL*, *CYBA*, *LYZ*, *CTSD*, *ITGB2*, *C3*, *CTSB*, *TYROBP*, *PSAP*, *PYCARD*, *CTSS*, *B2M*, *CD68*, *CD14*, *FCER1G* |
| Neutrophil Activation Involved In Immune Response | $2.53 \times 10^{-5}$ | *FTL*, *CYBA*, *LYZ*, *CTSD*, *ITGB2*, *C3*, *CTSB*, *TYROBP*, *PSAP*, *PYCARD*, *CTSS*, *B2M*, *CD68*, *CD14*, *FCER1G* |
| Regulation Of Cell Motility | $2.70 \times 10^{-5}$ | *APOE*, *CD74*, *SPARC*, *SFRP2*, *COL1A1*, *MDK*, *GPNMB*, *POSTN*, *FN1*, *PYCARD*, *SEMA3F*, *COL3A1*, *CORO1A*, *TIMP1*, *LGMN*, *EMILIN1*, *FLNA*, *DCN*, *SERPINF1*, *LRP1*, *RARRES2* |
| Regulation Of Multicellular Organismal Development | $2.98 \times 10^{-5}$ | *APOE*, *CD74*, *LGALS1*, *SPARC*, *C1QC*, *SFRP2*, *COL1A1*, *MDK*, *GPNMB*, *VIM*, *ITGB2*, *C3*, *CTHRC1*, *FN1*, *TYROBP*, *SEMA3F*, *COL3A1*, *TYMP*, *TIMP1*, *EMILIN1*, *FLNA*, *DCN*, *B2M*, *COMP*, *SERPINF1*, *INPP5J*, *LRP1*, *HSPG2*, *SPI1*, *PSMB8*, *FBN1* |
| Antigen Processing And Presentation | $3.03 \times 10^{-5}$ | *CD74*, *CYBA*, *CTSD*, *PYCARD*, *CTSS*, *LGMN*, *B2M*, *CD68*, *FCER1G*, *TAPBP*, *PSMB8* |
| Wound Healing | $3.33 \times 10^{-5}$ | *APOE*, *SPARC*, *COL1A1*, *MDK*, *A2M*, *POSTN*, *COL1A2*, *FN1*, *COL3A1*, *SERPING1*, *TIMP1*, *FLNA*, *DCN*, *COMP*, *FCER1G*, *ACTB* |
| Neutrophil Activation | $3.40 \times 10^{-5}$ | *FTL*, *CYBA*, *LYZ*, *CTSD*, *ITGB2*, *C3*, *CTSB*, *TYROBP*, *PSAP*, *PYCARD*, *CTSS*, *B2M*, *CD68*, *CD14*, *FCER1G* |
| Neutrophil Mediated Immunity | $3.40 \times 10^{-5}$ | *FTL*, *CYBA*, *LYZ*, *CTSD*, *ITGB2*, *C3*, *CTSB*, *TYROBP*, *PSAP*, *PYCARD*, *CTSS*, *B2M*, *CD68*, *CD14*, *FCER1G* |
| Granulocyte Activation | $3.99 \times 10^{-5}$ | *FTL*, *CYBA*, *LYZ*, *CTSD*, *ITGB2*, *C3*, *CTSB*, *TYROBP*, *PSAP*, *PYCARD*, *CTSS*, *B2M*, *CD68*, *CD14*, *FCER1G* |
| Multi-Organism Process | $4.21 \times 10^{-5}$ | *APOE*, *CD74*, *C1QB*, *C1QA*, *LGALS1*, *SPARC*, *C1QC*, *CYBA*, *MDK*, *A2M*, *LYZ*, *MUC1*, *VIM*, *ITGB2*, *C3*, *FN1*, *CTSB*, *TYROBP*, *PYCARD*, *FAU*, *RPLP1*, *SERPING1*, *CORO1A*, *TIMP1*, *CTSS*, *LGMN*, *FLNA*, *DCN*, *B2M*, *CD68*, *CD14*, *FCER1G*, *C1S*, *RPL10*, *RARRES2*, *RPL13*, *PSMB8* |
| Supramolecular Fiber Organization | $4.42 \times 10^{-5}$ | *APOE*, *CAPG*, *SFRP2*, *AEBP1*, *COL1A1*, *LUM*, *VIM*, *COL1A2*, *PYCARD*, *COL3A1*, *CORO1A*, *MMP11*, *EMILIN1*, *FLNA*, *B2M*, *COMP*, *INPP5J* |
| Positive Regulation Of Response To Stimulus | $5.98 \times 10^{-5}$ | *APOE*, *CD74*, *C1QB*, *C1QA*, *LGALS1*, *C1QC*, *CYBA*, *SFRP2*, *COL1A1*, *MDK*, *A2M*, *GPNMB*, *MUC1*, *ITGB2*, *C3*, *CTSB*, *PSAP*, *PYCARD*, *COL3A1*, *SERPING1*, *CTSS*, *LGMN*, *FLNA*, *DCN*, *B2M*, *CD14*, *FCER1G*, *C1S*, *LRP1*, *RARRES2*, *ACTB*, *PSMB8*, *TGM2* |
| Antigen Processing And Presentation Of Peptide Antigen | $6.05 \times 10^{-5}$ | *CD74*, *CYBA*, *CTSD*, *PYCARD*, *CTSS*, *LGMN*, *B2M*, *FCER1G*, *TAPBP*, *PSMB8* |
| Blood Vessel Morphogenesis | $6.37 \times 10^{-5}$ | *APOE*, *SPARC*, *SFRP2*, *MDK*, *GPNMB*, *ITGB2*, *C3*, *FN1*, *COL3A1*, *TYMP*, *EMILIN1*, *DCN*, *COMP*, *SERPINF1*, *LRP1*, *HSPG2*, *SPI1* |
| Skeletal System Development | $6.67 \times 10^{-5}$ | *SPARC*, *COL6A1*, *SFRP2*, *COL1A1*, *LUM*, *MDK*, *BGN*, *COL1A2*, *TYROBP*, *COL3A1*, *COL6A2*, *TIMP1*, *COMP*, *COL6A3*, *FBN1* |
| Regulation Of Locomotion | $8.66 \times 10^{-5}$ | *APOE*, *CD74*, *SPARC*, *SFRP2*, *COL1A1*, *MDK*, *GPNMB*, *POSTN*, *FN1*, *PYCARD*, *SEMA3F*, *COL3A1*, *CORO1A*, *TIMP1*, *LGMN*, *EMILIN1*, *FLNA*, *DCN*, *SERPINF1*, *LRP1*, *RARRES2* |
| Cell Surface Receptor Signaling Pathway | $8.99 \times 10^{-5}$ | *APOE*, *CD74*, *CYBA*, *SFRP2*, *COL1A1*, *MDK*, *REPS2*, *POSTN*, *MUC1*, *VIM*, *ITGB2*, *LY6E*, *C3*, *CTHRC1*, *COL1A2*, *FN1*, *TYROBP*, *PYCARD*, *SEMA3F*, *COL3A1*, *CORO1A*, *TIMP1*, *LGMN*, *EMILIN1*, *FLNA*, *GFRA1*, *DCN*, *B2M*, *COMP*, *CD14*, *FCER1G*, *LRP1*, *RARRES2*, *ACTB*, *SPI1*, *PSMB8*, *FBN1* |
| Myeloid Cell Activation Involved In Immune Response | $1.01 \times 10^{-4}$ | *FTL*, *CYBA*, *LYZ*, *CTSD*, *ITGB2*, *C3*, *CTSB*, *TYROBP*, *PSAP*, *PYCARD*, *CTSS*, *B2M*, *CD68*, *CD14*, *FCER1G* |
| Response To Stimulus | $1.13 \times 10^{-4}$ | *APOE*, *CD74*, *C1QB*, *FTL*, *C1QA*, *LGALS1*, *SPARC*, *C1QC*, *CRIP1*, *COL6A1*, *LAPTM5*, *CYBA*, *CPNE7*, *SFRP2*, *COL1A1*, *LUM*, *MDK*, *A2M*, *REPS2*, *GPNMB*, *LYZ*, *POSTN*, *MUC1*, *CTSD*, *VIM*, *ITGB2*, *LY6E*, *C3*, *CTHRC1*, *COL1A2*, *FN1*, *CTSB*, *TYROBP*, *PSAP*, *PYCARD*, *FAU*, *SELENOM*, *SEMA3F*, *COL3A1*, *COL6A2*, *TYMP*, *SERPING1*, *CORO1A*, *TIMP1*, *CTSS*, *LGMN*, *EMILIN1*, *FLNA*, *GFRA1*, *DCN*, *B2M*, *COMP*, *CD68*, *CD14*, *SERPINF1*, *FCER1G*, *C1S*, *LRP1*, *RARRES2*, *ACTB*, *TAPBP*, *ELOB*, *VSTM2A*, *HSPG2*, *SPI1*, *SPARCL1*, *LSP1*, *PSMB8*, *TGM2*, *FBN1* |
| Regulation Of Cellular Component Movement | $1.13 \times 10^{-4}$ | *APOE*, *CD74*, *SPARC*, *SFRP2*, *COL1A1*, *MDK*, *GPNMB*, *POSTN*, *FN1*, *PYCARD*, *SEMA3F*, *COL3A1*, *CORO1A*, *TIMP1*, *LGMN*, *EMILIN1*, *FLNA*, *DCN*, *SERPINF1*, *LRP1*, *RARRES2* |

| Biological process | Adjusted *p*-value | Intersecting genes |
|---|---|---|
| Myeloid Leukocyte Mediated Immunity | $1.20 \times 10^{-4}$ | *FTL*, *CYBA*, *LYZ*, *CTSD*, *ITGB2*, *C3*, *CTSB*, *TYROBP*, *PSAP*, *PYCARD*, *CTSS*, *B2M*, *CD68*, *CD14*, *FCER1G* |
| Anatomical Structure Development | $1.26 \times 10^{-4}$ | *APOE*, *CD74*, *C1QB*, *C1QA*, *LGALS1*, *SPARC*, *C1QC*, *CRIP1*, *COL6A1*, *SFRP2*, *CRIP2*, *COL1A1*, *LUM*, *MDK*, *GPNMB*, *POSTN*, *VIM*, *SLCO2B1*, *ITGB2*, *C3*, *BGN*, *CTHRC1*, *COL1A2*, *FN1*, *CTSB*, *TYROBP*, *PSAP*, *SELENOM*, *SEMA3F*, *COL3A1*, *COL6A2*, *TYMP*, *CORO1A*, *TIMP1*, *MMP11*, *EMILIN1*, *FLNA*, *GFRA1*, *DCN*, *B2M*, *COMP*, *SERPINF1*, *INPP5J*, *FCER1G*, *RPL10*, *LRP1*, *RARRES2*, *ACTB*, *COL6A3*, *HSPG2*, *SPI1*, *PSMB8*, *TGM2*, *FBN1* |
| Regulation Of Response To External Stimulus | $1.42 \times 10^{-4}$ | *APOE*, *CD74*, *CYBA*, *MDK*, *A2M*, *MUC1*, *ITGB2*, *C3*, *CTSB*, *PYCARD*, *SEMA3F*, *SERPING1*, *CTSS*, *LGMN*, *CD14*, *SERPINF1*, *FCER1G*, *LRP1*, *RARRES2*, *PSMB8*, *TGM2* |
| Positive Regulation Of Response To External Stimulus | $1.44 \times 10^{-4}$ | *CD74*, *CYBA*, *MDK*, *MUC1*, *ITGB2*, *C3*, *CTSB*, *PYCARD*, *CTSS*, *LGMN*, *CD14*, *FCER1G*, *LRP1*, *RARRES2*, *PSMB8*, *TGM2* |
| Anatomical Structure Morphogenesis | $1.69 \times 10^{-4}$ | *APOE*, *SPARC*, *CRIP1*, *COL6A1*, *SFRP2*, *COL1A1*, *MDK*, *GPNMB*, *POSTN*, *ITGB2*, *C3*, *CTHRC1*, *COL1A2*, *FN1*, *TYROBP*, *SEMA3F*, *COL3A1*, *COL6A2*, *TYMP*, *CORO1A*, *EMILIN1*, *FLNA*, *GFRA1*, *DCN*, *COMP*, *SERPINF1*, *LRP1*, *ACTB*, *COL6A3*, *HSPG2*, *SPI1*, *PSMB8*, *TGM2*, *FBN1* |
| Cartilage Development | $1.78 \times 10^{-4}$ | *COL6A1*, *SFRP2*, *COL1A1*, *LUM*, *MDK*, *BGN*, *COL6A2*, *TIMP1*, *COMP*, *COL6A3* |
| Collagen Metabolic Process | $1.85 \times 10^{-4}$ | *COL1A1*, *CTSD*, *VIM*, *COL1A2*, *CTSB*, *MMP11*, *CTSS*, *EMILIN1* |
| Tube Morphogenesis | $1.87 \times 10^{-4}$ | *APOE*, *SPARC*, *SFRP2*, *MDK*, *GPNMB*, *ITGB2*, *C3*, *CTHRC1*, *FN1*, *COL3A1*, *TYMP*, *EMILIN1*, *DCN*, *COMP*, *SERPINF1*, *LRP1*, *HSPG2*, *SPI1*, *TGM2* |
| Cell Migration | $1.97 \times 10^{-4}$ | *APOE*, *CD74*, *SPARC*, *SFRP2*, *COL1A1*, *MDK*, *GPNMB*, *POSTN*, *ITGB2*, *CTHRC1*, *COL1A2*, *FN1*, *PYCARD*, *SEMA3F*, *COL3A1*, *CORO1A*, *TIMP1*, *LGMN*, *EMILIN1*, *FLNA*, *DCN*, *SERPINF1*, *FCER1G*, *LRP1*, *RARRES2* |
| Locomotion | $2.31 \times 10^{-4}$ | *APOE*, *CD74*, *SPARC*, *SFRP2*, *COL1A1*, *MDK*, *GPNMB*, *POSTN*, *ITGB2*, *CTHRC1*, *COL1A2*, *FN1*, *PYCARD*, *SEMA3F*, *COL3A1*, *TYMP*, *CORO1A*, *TIMP1*, *LGMN*, *EMILIN1*, *FLNA*, *GFRA1*, *DCN*, *SERPINF1*, *FCER1G*, *LRP1*, *RARRES2*, *ACTB* |
| Bone Development | $2.41 \times 10^{-4}$ | *SPARC*, *COL6A1*, *SFRP2*, *COL1A1*, *BGN*, *TYROBP*, *COL6A2*, *COMP*, *COL6A3*, *FBN1* |
| Regulation Of Response To Stimulus | $2.68 \times 10^{-4}$ | *APOE*, *CD74*, *C1QB*, *C1QA*, *LGALS1*, *C1QC*, *CYBA*, *SFRP2*, *COL1A1*, *MDK*, *A2M*, *GPNMB*, *POSTN*, *MUC1*, *ITGB2*, *LY6E*, *C3*, *CTHRC1*, *COL1A2*, *FN1*, *CTSB*, *TYROBP*, *PSAP*, *PYCARD*, *SEMA3F*, *COL3A1*, *SERPING1*, *TIMP1*, *CTSS*, *LGMN*, *EMILIN1*, *FLNA*, *DCN*, *B2M*, *CD14*, *SERPINF1*, *FCER1G*, *C1S*, *LRP1*, *RARRES2*, *ACTB*, *PSMB8*, *TGM2*, *FBN1* |
| Negative Regulation Of Biological Process | $2.76 \times 10^{-4}$ | *APOE*, *CD74*, *APOC1*, *CAPG*, *LGALS1*, *SPARC*, *C1QC*, *CYBA*, *SFRP2*, *AEBP1*, *COL1A1*, *MDK*, *A2M*, *GPNMB*, *POSTN*, *MUC1*, *VIM*, *ITGB2*, *C3*, *CTHRC1*, *FN1*, *TYROBP*, *PSAP*, *PYCARD*, *FAU*, *RPLP1*, *SEMA3F*, *COL3A1*, *SERPING1*, *CORO1A*, *TIMP1*, *MMP11*, *LGMN*, *EMILIN1*, *FLNA*, *DCN*, *B2M*, *COMP*, *CD68*, *CD14*, *SERPINF1*, *INPP5J*, *FCER1G*, *RPL10*, *LRP1*, *RPL13*, *COL6A3*, *HSPG2*, *SPI1*, *PSMB8*, *FBN1* |
| Cell Motility | $2.92 \times 10^{-4}$ | *APOE*, *CD74*, *SPARC*, *SFRP2*, *COL1A1*, *MDK*, *GPNMB*, *POSTN*, *ITGB2*, *CTHRC1*, *COL1A2*, *FN1*, *PYCARD*, *SEMA3F*, *COL3A1*, *CORO1A*, *TIMP1*, *LGMN*, *EMILIN1*, *FLNA*, *DCN*, *SERPINF1*, *FCER1G*, *LRP1*, *RARRES2*, *ACTB* |
| Localization Of Cell | $2.92 \times 10^{-4}$ | *APOE*, *CD74*, *SPARC*, *SFRP2*, *COL1A1*, *MDK*, *GPNMB*, *POSTN*, *ITGB2*, *CTHRC1*, *COL1A2*, *FN1*, *PYCARD*, *SEMA3F*, *COL3A1*, *CORO1A*, *TIMP1*, *LGMN*, *EMILIN1*, *FLNA*, *DCN*, *SERPINF1*, *FCER1G*, *LRP1*, *RARRES2*, *ACTB* |
| Multicellular Organismal Process | $3.13 \times 10^{-4}$ | *APOE*, *CD74*, *APOC1*, *C1QB*, *C1QA*, *LGALS1*, *SPARC*, *C1QC*, *CRIP1*, *COL6A1*, *CYBA*, *SFRP2*, *CRIP2*, *COL1A1*, *LUM*, *MDK*, *A2M*, *GPNMB*, *LYZ*, *POSTN*, *VIM*, *SLCO2B1*, *ITGB2*, *C3*, *BGN*, *CTHRC1*, *COL1A2*, *FN1*, *CTSB*, *TYROBP*, *PSAP*, *PYCARD*, *SELENOM*, *SEMA3F*, *COL3A1*, *COL6A2*, *TYMP*, *SERPING1*, *CORO1A*, *TIMP1*, *MMP11*, *LGMN*, *EMILIN1*, *FLNA*, *GFRA1*, *DCN*, *B2M*, *COMP*, *CD14*, *SERPINF1*, *INPP5J*, *FCER1G*, *RPL10*, *LRP1*, *RARRES2*, *ACTB*, *COL6A3*, *HSPG2*, *SPI1*, *PSMB8*, *TGM2*, *FBN1* |
| Antigen Processing And Presentation Of Exogenous Peptide Antigen | $3.26 \times 10^{-4}$ | *CD74*, *CYBA*, *CTSD*, *CTSS*, *LGMN*, *B2M*, *FCER1G*, *TAPBP*, *PSMB8* |
| Circulatory System Development | $3.38 \times 10^{-4}$ | *APOE*, *SPARC*, *CRIP1*, *SFRP2*, *COL1A1*, *MDK*, *GPNMB*, *ITGB2*, *C3*, *COL1A2*, *FN1*, *COL3A1*, *TYMP*, *EMILIN1*, *DCN*, *COMP*, *SERPINF1*, *LRP1*, *HSPG2*, *SPI1*, *FBN1* |
| Synapse Pruning | $3.53 \times 10^{-4}$ | *C1QB*, *C1QA*, *C1QC*, *C3* |

| **Biological process** | **Adjusted *p*-value** | **Intersecting genes** |
|---|---|---|
| Leukocyte Activation | $3.58 \times 10^{-4}$ | *CD74*, *FTL*, *C1QA*, *LGALS1*, *CYBA*, *MDK*, *GPNMB*, *LYZ*, *CTSD*, *ITGB2*, *C3*, *CTSB*, *TYROBP*, *PSAP*, *PYCARD*, *CORO1A*, *CTSS*, *B2M*, *CD68*, *CD14*, *FCER1G*, *SPI1* |
| Regulation Of Cell Adhesion | $4.50 \times 10^{-4}$ | *CD74*, *LGALS1*, *SFRP2*, *COL1A1*, *MDK*, *GPNMB*, *POSTN*, *MUC1*, *ITGB2*, *FN1*, *PYCARD*, *CORO1A*, *EMILIN1*, *FLNA*, *LRP1*, *TGM2* |
| Antigen Processing And Presentation Of Exogenous Antigen | $4.58 \times 10^{-4}$ | *CD74*, *CYBA*, *CTSD*, *CTSS*, *LGMN*, *B2M*, *FCER1G*, *TAPBP*, *PSMB8* |
| Regulation Of Innate Immune Response | $5.34 \times 10^{-4}$ | *APOE*, *CYBA*, *A2M*, *MUC1*, *ITGB2*, *CTSB*, *PYCARD*, *SERPING1*, *CTSS*, *LGMN*, *CD14*, *FCER1G*, *PSMB8* |
| Regulation Of Defense Response | $5.72 \times 10^{-4}$ | *APOE*, *CYBA*, *MDK*, *A2M*, *MUC1*, *ITGB2*, *C3*, *CTSB*, *PYCARD*, *SERPING1*, *CTSS*, *LGMN*, *CD14*, *SERPINF1*, *FCER1G*, *PSMB8*, *TGM2* |
| Protein Metabolic Process | $5.95 \times 10^{-4}$ | *APOE*, *CD74*, *APOC1*, *LGALS1*, *SFRP2*, *AEBP1*, *A2M*, *GPNMB*, *LYZ*, *MUC1*, *CTSD*, *VIM*, *ITGB2*, *C3*, *BGN*, *FN1*, *CTSB*, *TYROBP*, *PSAP*, *PYCARD*, *FAU*, *RPLP1*, *COL3A1*, *SERPING1*, *TIMP1*, *MMP11*, *CTSS*, *LGMN*, *EMILIN1*, *FLNA*, *GFRA1*, *DCN*, *CPB1*, *B2M*, *COMP*, *GALNT6*, *SERPINF1*, *INPP5J*, *C1S*, *RPL10*, *LRP1*, *RARRES2*, *ACTB*, *RPL13*, *ELOB*, *COL6A3*, *HSPG2*, *SPI1*, *SPARCL1*, *PSMB8*, *TGM2*, *FBN1* |
| Tissue Development | $6.30 \times 10^{-4}$ | *CRIP1*, *COL6A1*, *SFRP2*, *COL1A1*, *LUM*, *MDK*, *GPNMB*, *POSTN*, *VIM*, *ITGB2*, *BGN*, *CTHRC1*, *COL1A2*, *FN1*, *CTSB*, *PSAP*, *SELENOM*, *SEMA3F*, *COL3A1*, *COL6A2*, *TIMP1*, *FLNA*, *DCN*, *COMP*, *COL6A3*, *HSPG2*, *PSMB8*, *TGM2* |
| Regulation Of Peptidase Activity | $6.37 \times 10^{-4}$ | *SFRP2*, *A2M*, *CTSD*, *C3*, *FN1*, *PYCARD*, *SERPING1*, *TIMP1*, *LGMN*, *SERPINF1*, *LRP1*, *COL6A3*, *PSMB8* |
| Tube Development | $7.33 \times 10^{-4}$ | *APOE*, *SPARC*, *SFRP2*, *MDK*, *GPNMB*, *ITGB2*, *C3*, *CTHRC1*, *FN1*, *COL3A1*, *TYMP*, *EMILIN1*, *DCN*, *COMP*, *SERPINF1*, *LRP1*, *RARRES2*, *HSPG2*, *SPI1*, *TGM2* |
| Positive Regulation Of Cell Migration | $7.72 \times 10^{-4}$ | *CD74*, *SPARC*, *COL1A1*, *MDK*, *GPNMB*, *POSTN*, *FN1*, *PYCARD*, *SEMA3F*, *CORO1A*, *LGMN*, *FLNA*, *LRP1*, *RARRES2* |
| Response To Oxygen-Containing Compound | $8.07 \times 10^{-4}$ | *APOE*, *LGALS1*, *SPARC*, *COL6A1*, *CYBA*, *COL1A1*, *POSTN*, *VIM*, *COL1A2*, *PSAP*, *PYCARD*, *COL3A1*, *COL6A2*, *TIMP1*, *LGMN*, *FLNA*, *DCN*, *CD68*, *CD14*, *SERPINF1*, *LRP1*, *RARRES2*, *ACTB*, *SPI1*, *FBN1* |
| Negative Regulation Of Cellular Process | $1.04 \times 10^{-3}$ | *APOE*, *CD74*, *APOC1*, *CAPG*, *LGALS1*, *SPARC*, *C1QC*, *CYBA*, *SFRP2*, *AEBP1*, *COL1A1*, *MDK*, *A2M*, *GPNMB*, *POSTN*, *MUC1*, *VIM*, *ITGB2*, *C3*, *CTHRC1*, *FN1*, *TYROBP*, *PSAP*, *PYCARD*, *SEMA3F*, *COL3A1*, *SERPING1*, *CORO1A*, *TIMP1*, *MMP11*, *LGMN*, *EMILIN1*, *FLNA*, *DCN*, *B2M*, *COMP*, *CD14*, *SERPINF1*, *INPP5J*, *FCER1G*, *RPL10*, *LRP1*, *COL6A3*, *SPI1*, *PSMB8*, *FBN1* |
| Positive Regulation Of Cell Motility | $1.24 \times 10^{-3}$ | *CD74*, *SPARC*, *COL1A1*, *MDK*, *GPNMB*, *POSTN*, *FN1*, *PYCARD*, *SEMA3F*, *CORO1A*, *LGMN*, *FLNA*, *LRP1*, *RARRES2* |
| Negative Regulation Of Immune System Process | $1.28 \times 10^{-3}$ | *CD74*, *C1QC*, *MDK*, *A2M*, *GPNMB*, *TYROBP*, *COL3A1*, *SERPING1*, *EMILIN1*, *CD68*, *CD14*, *FCER1G*, *FBN1* |
| Regulation Of Anatomical Structure Morphogenesis | $1.33 \times 10^{-3}$ | *APOE*, *SPARC*, *SFRP2*, *MDK*, *GPNMB*, *POSTN*, *ITGB2*, *C3*, *CTHRC1*, *FN1*, *TYROBP*, *SEMA3F*, *CORO1A*, *EMILIN1*, *FLNA*, *DCN*, *SERPINF1*, *LRP1*, *HSPG2*, *PSMB8* |
| Localization | $1.42 \times 10^{-3}$ | *APOE*, *CD74*, *APOC1*, *FTL*, *SPARC*, *CYBA*, *SFRP2*, *COL1A1*, *MDK*, *A2M*, *GPNMB*, *LYZ*, *POSTN*, *CTSD*, *SLCO2B1*, *ITGB2*, *ISLR*, *C3*, *CTHRC1*, *COL1A2*, *FN1*, *CTSB*, *TYROBP*, *PSAP*, *PYCARD*, *FAU*, *RPLP1*, *SELENOM*, *SEMA3F*, *COL3A1*, *SERPING1*, *CORO1A*, *TIMP1*, *CTSS*, *LGMN*, *EMILIN1*, *FLNA*, *DCN*, *B2M*, *SLC39A6*, *COMP*, *CD68*, *CD14*, *SERPINF1*, *FCER1G*, *RPL10*, *LRP1*, *RARRES2*, *ACTB*, *TAPBP*, *RPL13*, *VSTM2A*, *HSPG2*, *PSMB8*, *TGM2*, *FBN1* |
| Peptide Cross-Linking | $1.48 \times 10^{-3}$ | *BGN*, *FN1*, *COL3A1*, *DCN*, *TGM2* |
| Cellular Developmental Process | $1.65 \times 10^{-3}$ | *APOE*, *CD74*, *C1QA*, *LGALS1*, *SPARC*, *C1QC*, *COL6A1*, *SFRP2*, *COL1A1*, *MDK*, *A2M*, *GPNMB*, *POSTN*, *VIM*, *ITGB2*, *C3*, *CTHRC1*, *FN1*, *CTSB*, *TYROBP*, *PSAP*, *SEMA3F*, *COL3A1*, *COL6A2*, *TYMP*, *CORO1A*, *MMP11*, *FLNA*, *GFRA1*, *B2M*, *COMP*, *SERPINF1*, *INPP5J*, *FCER1G*, *LRP1*, *RARRES2*, *ACTB*, *COL6A3*, *VSTM2A*, *HSPG2*, *SPI1*, *PSMB8*, *FBN1* |
| Regulation Of Localization | $1.74 \times 10^{-3}$ | *APOE*, *CD74*, *APOC1*, *SPARC*, *CYBA*, *SFRP2*, *COL1A1*, *MDK*, *GPNMB*, *POSTN*, *CTSD*, *ITGB2*, *C3*, *FN1*, *TYROBP*, *PYCARD*, *SEMA3F*, *COL3A1*, *CORO1A*, *TIMP1*, *CTSS*, *LGMN*, *EMILIN1*, *FLNA*, *DCN*, *B2M*, *CD14*, *SERPINF1*, *FCER1G*, *LRP1*, *RARRES2*, *ACTB*, *VSTM2A* |
| Positive Regulation Of Cellular Component Movement | $1.80 \times 10^{-3}$ | *CD74*, *SPARC*, *COL1A1*, *MDK*, *GPNMB*, *POSTN*, *FN1*, *PYCARD*, *SEMA3F*, *CORO1A*, *LGMN*, *FLNA*, *LRP1*, *RARRES2* |
| Regulation Of Multi-Organism Process | $1.82 \times 10^{-3}$ | *APOE*, *CD74*, *LGALS1*, *CYBA*, *A2M*, *MUC1*, *ITGB2*, *CTSB*, *PYCARD*, *SERPING1*, *TIMP1*, *CTSS*, *LGMN*, *CD14*, *FCER1G*, *PSMB8* |

| **Biological process** | **Adjusted *p*-value** | **Intersecting genes** |
|---|---|---|
| Platelet Activation | $1.84 \times 10^{-3}$ | *APOE*, *COL1A1*, *COL1A2*, *COL3A1*, *FLNA*, *COMP*, *FCER1G*, *ACTB* |
| Positive Regulation Of Locomotion | $2.20 \times 10^{-3}$ | *CD74*, *SPARC*, *COL1A1*, *MDK*, *GPNMB*, *POSTN*, *FN1*, *PYCARD*, *SEMA3F*, *CORO1A*, *LGMN*, *FLNA*, *LRP1*, *RARRES2* |
| Cell-Substrate Adhesion | $2.24 \times 10^{-3}$ | *LGALS1*, *COL1A1*, *MDK*, *POSTN*, *ITGB2*, *FN1*, *COL3A1*, *CORO1A*, *EMILIN1*, *FLNA*, *LRP1* |
| Regulation Of Complement Activation | $2.26 \times 10^{-3}$ | *C1QB*, *C1QA*, *C1QC*, *A2M*, *C3*, *SERPING1*, *C1S* |
| Positive Regulation Of Defense Response | $2.30 \times 10^{-3}$ | *CYBA*, *MDK*, *MUC1*, *ITGB2*, *C3*, *CTSB*, *PYCARD*, *CTSS*, *LGMN*, *CD14*, *FCER1G*, *PSMB8*, *TGM2* |
| Movement Of Cell Or Subcellular Component | $2.42 \times 10^{-3}$ | *APOE*, *CD74*, *SPARC*, *SFRP2*, *COL1A1*, *MDK*, *GPNMB*, *POSTN*, *VIM*, *ITGB2*, *CTHRC1*, *COL1A2*, *FN1*, *PYCARD*, *SEMA3F*, *COL3A1*, *CORO1A*, *TIMP1*, *LGMN*, *EMILIN1*, *FLNA*, *GFRA1*, *DCN*, *SERPINF1*, *FCER1G*, *LRP1*, *RARRES2*, *ACTB* |
| Response To Cytokine | $2.46 \times 10^{-3}$ | *CD74*, *SPARC*, *LAPTM5*, *CYBA*, *COL1A1*, *POSTN*, *MUC1*, *VIM*, *ITGB2*, *COL1A2*, *FN1*, *PYCARD*, *COL3A1*, *CORO1A*, *TIMP1*, *B2M*, *CD14*, *FCER1G*, *SPI1*, *PSMB8* |
| Regulation Of Response To Biotic Stimulus | $2.79 \times 10^{-3}$ | *APOE*, *CYBA*, *A2M*, *MUC1*, *ITGB2*, *CTSB*, *PYCARD*, *SERPING1*, *CTSS*, *LGMN*, *CD14*, *FCER1G*, *PSMB8* |
| Regulation Of Immune Effector Process | $3.63 \times 10^{-3}$ | *CD74*, *C1QB*, *C1QA*, *C1QC*, *A2M*, *ITGB2*, *C3*, *PYCARD*, *SERPING1*, *B2M*, *FCER1G*, *C1S* |
| Positive Regulation Of Multi-Organism Process | $3.83 \times 10^{-3}$ | *APOE*, *CD74*, *LGALS1*, *CYBA*, *MUC1*, *ITGB2*, *CTSB*, *PYCARD*, *CTSS*, *LGMN*, *CD14*, *FCER1G*, *PSMB8* |
| Cell Differentiation | $4.21 \times 10^{-3}$ | *APOE*, *CD74*, *C1QA*, *LGALS1*, *C1QC*, *COL6A1*, *SFRP2*, *COL1A1*, *MDK*, *A2M*, *GPNMB*, *POSTN*, *VIM*, *ITGB2*, *C3*, *CTHRC1*, *FN1*, *CTSB*, *TYROBP*, *PSAP*, *SEMA3F*, *COL3A1*, *COL6A2*, *TYMP*, *MMP11*, *FLNA*, *GFRA1*, *B2M*, *COMP*, *SERPINF1*, *INPP5J*, *FCER1G*, *LRP1*, *RARRES2*, *ACTB*, *COL6A3*, *VSTM2A*, *HSPG2*, *SPI1*, *PSMB8*, *FBN1* |
| Phagocytosis | $4.26 \times 10^{-3}$ | *CYBA*, *ITGB2*, *C3*, *TYROBP*, *PYCARD*, *CORO1A*, *CD14*, *FCER1G*, *LRP1*, *ACTB*, *TGM2* |
| Regulation Of Cell Death | $4.70 \times 10^{-3}$ | *APOE*, *CD74*, *C1QA*, *LGALS1*, *SFRP2*, *MDK*, *GPNMB*, *MUC1*, *CTSD*, *ITGB2*, *CTSB*, *TYROBP*, *PSAP*, *PYCARD*, *CORO1A*, *TIMP1*, *LGMN*, *FLNA*, *COMP*, *SERPINF1*, *FCER1G*, *RPL10*, *LRP1*, *TGM2* |
| Organonitrogen Compound Metabolic Process | $4.86 \times 10^{-3}$ | *APOE*, *CD74*, *APOC1*, *LGALS1*, *SFRP2*, *AEBP1*, *LUM*, *A2M*, *GPNMB*, *LYZ*, *MUC1*, *CTSD*, *VIM*, *ITGB2*, *C3*, *BGN*, *FN1*, *CTSB*, *TYROBP*, *PSAP*, *PYCARD*, *FAU*, *RPLP1*, *COL3A1*, *TYMP*, *SERPING1*, *TIMP1*, *MMP11*, *CTSS*, *LGMN*, *EMILIN1*, *FLNA*, *GFRA1*, *DCN*, *CPB1*, *B2M*, *COMP*, *GALNT6*, *SERPINF1*, *INPP5J*, *C1S*, *RPL10*, *LRP1*, *RARRES2*, *ACTB*, *TAPBP*, *RPL13*, *ELOB*, *COL6A3*, *HSPG2*, *SPI1*, *SPARCL1*, *PSMB8*, *TGM2*, *FBN1* |
| Regulation Of Cell Differentiation | $5.32 \times 10^{-3}$ | *APOE*, *CD74*, *LGALS1*, *C1QC*, *SFRP2*, *COL1A1*, *MDK*, *POSTN*, *VIM*, *CTHRC1*, *FN1*, *TYROBP*, *SEMA3F*, *COL3A1*, *MMP11*, *FLNA*, *B2M*, *SERPINF1*, *INPP5J*, *LRP1*, *RARRES2*, *VSTM2A*, *SPI1*, *PSMB8*, *FBN1* |
| Positive Regulation Of Biological Process | $5.35 \times 10^{-3}$ | *APOE*, *CD74*, *APOC1*, *C1QB*, *C1QA*, *LGALS1*, *SPARC*, *C1QC*, *CYBA*, *SFRP2*, *CRIP2*, *COL1A1*, *LUM*, *MDK*, *A2M*, *GPNMB*, *POSTN*, *MUC1*, *CTSD*, *VIM*, *ITGB2*, *C3*, *CTHRC1*, *FN1*, *CTSB*, *TYROBP*, *PSAP*, *PYCARD*, *RPLP1*, *SEMA3F*, *COL3A1*, *SERPING1*, *CORO1A*, *TIMP1*, *CTSS*, *LGMN*, *EMILIN1*, *FLNA*, *DCN*, *B2M*, *COMP*, *CD14*, *SERPINF1*, *FCER1G*, *C1S*, *LRP1*, *RARRES2*, *ACTB*, *VSTM2A*, *SPI1*, *PSMB8*, *TGM2* |
| Chondrocyte Development | $5.98 \times 10^{-3}$ | *COL6A1*, *SFRP2*, *COL6A2*, *COMP*, *COL6A3* |
| Positive Regulation Of Cytokine Production | $6.10 \times 10^{-3}$ | *CD74*, *CYBA*, *LUM*, *MDK*, *POSTN*, *C3*, *TYROBP*, *PYCARD*, *B2M*, *CD14*, *FCER1G*, *LRP1* |
| Response To Organonitrogen Compound | $6.49 \times 10^{-3}$ | *APOE*, *SPARC*, *COL6A1*, *CYBA*, *COL1A1*, *VIM*, *ITGB2*, *COL1A2*, *COL3A1*, *TIMP1*, *LGMN*, *FLNA*, *CD68*, *FCER1G*, *LRP1*, *RARRES2*, *ACTB*, *FBN1* |
| Regulation Of Humoral Immune Response | $6.59 \times 10^{-3}$ | *C1QB*, *C1QA*, *C1QC*, *A2M*, *C3*, *SERPING1*, *C1S* |
| Cartilage Development Involved In Endochondral Bone Morphogenesis | $6.67 \times 10^{-3}$ | *COL6A1*, *COL1A1*, *COL6A2*, *COMP*, *COL6A3* |
| Receptor-Mediated Endocytosis | $7.13 \times 10^{-3}$ | *APOE*, *APOC1*, *SPARC*, *ITGB2*, *C3*, *B2M*, *CD14*, *FCER1G*, *LRP1*, *HSPG2* |
| Negative Regulation Of Cell Differentiation | $7.40 \times 10^{-3}$ | *APOE*, *CD74*, *LGALS1*, *C1QC*, *SFRP2*, *MDK*, *POSTN*, *VIM*, *SEMA3F*, *COL3A1*, *MMP11*, *B2M*, *INPP5J*, *LRP1*, *FBN1* |
| Regulation Of Response To Stress | $7.65 \times 10^{-3}$ | *APOE*, *CD74*, *CYBA*, *SFRP2*, *MDK*, *A2M*, *MUC1*, *ITGB2*, *C3*, *CTSB*, *PSAP*, *PYCARD*, *SERPING1*, *CTSS*, *LGMN*, *B2M*, *CD14*, *SERPINF1*, *FCER1G*, *LRP1*, *PSMB8*, *TGM2* |

| Biological process | Adjusted *p*-value | Intersecting genes |
|---|---|---|
| Activation Of Innate Immune Response | $7.75 \times 10^{-3}$ | *CYBA*, *MUC1*, *ITGB2*, *CTSB*, *PYCARD*, *CTSS*, *LGMN*, *CD14*, *FCER1G*, *PSMB8* |
| Regulation Of Neuron Death | $8.19 \times 10^{-3}$ | *APOE*, *C1QA*, *MDK*, *GPNMB*, *ITGB2*, *TYROBP*, *CORO1A*, *LGMN*, *SERPINF1*, *LRP1* |
| Aging | $8.42 \times 10^{-3}$ | *C1QA*, *ITGB2*, *SERPING1*, *TIMP1*, *DCN*, *B2M*, *COMP*, *CD68*, *SERPINF1*, *LRP1* |
| Positive Regulation Of Cell Adhesion | $8.50 \times 10^{-3}$ | *CD74*, *LGALS1*, *SFRP2*, *MDK*, *ITGB2*, *FN1*, *PYCARD*, *CORO1A*, *EMILIN1*, *FLNA*, *TGM2* |
| Cellular Response To Oxygen-Containing Compound | $1.03 \times 10^{-2}$ | *LGALS1*, *COL6A1*, *CYBA*, *COL1A1*, *VIM*, *COL1A2*, *PSAP*, *PYCARD*, *COL3A1*, *LGMN*, *FLNA*, *CD68*, *CD14*, *SERPINF1*, *LRP1*, *RARRES2*, *ACTB*, *SPI1*, *FBN1* |
| Leukocyte Migration | $1.08 \times 10^{-2}$ | *CD74*, *COL1A1*, *MDK*, *ITGB2*, *COL1A2*, *FN1*, *PYCARD*, *CORO1A*, *LGMN*, *EMILIN1*, *FCER1G*, *RARRES2* |
| Cellular Response To Endogenous Stimulus | $1.11 \times 10^{-2}$ | *COL6A1*, *CYBA*, *SFRP2*, *COL1A1*, *POSTN*, *VIM*, *COL1A2*, *CTSB*, *COL3A1*, *CORO1A*, *CTSS*, *LGMN*, *EMILIN1*, *FLNA*, *COMP*, *SERPINF1*, *LRP1*, *RARRES2*, *ACTB*, *VSTM2A*, *FBN1* |
| Regulation Of Biological Quality | $1.14 \times 10^{-2}$ | *APOE*, *CD74*, *FTL*, *CAPG*, *LGALS1*, *SPARC*, *CYBA*, *COL1A1*, *A2M*, *LYZ*, *POSTN*, *VIM*, *ITGB2*, *C3*, *COL1A2*, *FN1*, *TYROBP*, *PYCARD*, *SELENOM*, *SEMA3F*, *COL3A1*, *SERPING1*, *CORO1A*, *LGMN*, *FLNA*, *DCN*, *B2M*, *SLC39A6*, *COMP*, *SERPINF1*, *FCER1G*, *LRP1*, *ACTB*, *TAPBP*, *VSTM2A*, *SPI1*, *PSMB8*, *TGM2*, *FBN1* |
| Developmental Growth | $1.17 \times 10^{-2}$ | *APOE*, *COL6A1*, *SFRP2*, *MDK*, *POSTN*, *FN1*, *PSAP*, *SELENOM*, *SEMA3F*, *COL6A2*, *LGMN*, *COMP*, *LRP1*, *COL6A3* |
| Blood Coagulation | $1.22 \times 10^{-2}$ | *APOE*, *COL1A1*, *A2M*, *COL1A2*, *COL3A1*, *SERPING1*, *FLNA*, *COMP*, *FCER1G*, *ACTB* |
| Hemostasis | $1.39 \times 10^{-2}$ | *APOE*, *COL1A1*, *A2M*, *COL1A2*, *COL3A1*, *SERPING1*, *FLNA*, *COMP*, *FCER1G*, *ACTB* |
| Cell Death | $1.46 \times 10^{-2}$ | *APOE*, *CD74*, *C1QA*, *LGALS1*, *CRIP1*, *SFRP2*, *MDK*, *GPNMB*, *MUC1*, *CTSD*, *ITGB2*, *CTSB*, *TYROBP*, *PSAP*, *PYCARD*, *CORO1A*, *TIMP1*, *LGMN*, *FLNA*, *COMP*, *CD14*, *SERPINF1*, *FCER1G*, *RPL10*, *LRP1*, *SPI1*, *TGM2* |
| Negative Regulation Of Cell Development | $1.46 \times 10^{-2}$ | *APOE*, *LGALS1*, *POSTN*, *VIM*, *SEMA3F*, *COL3A1*, *B2M*, *INPP5J*, *LRP1*, *FBN1* |
| Coagulation | $1.50 \times 10^{-2}$ | *APOE*, *COL1A1*, *A2M*, *COL1A2*, *COL3A1*, *SERPING1*, *FLNA*, *COMP*, *FCER1G*, *ACTB* |
| Regulation Of Endopeptidase Activity | $1.54 \times 10^{-2}$ | *SFRP2*, *A2M*, *CTSD*, *C3*, *PYCARD*, *SERPING1*, *TIMP1*, *LGMN*, *SERPINF1*, *COL6A3*, *PSMB8* |
| Antigen Processing And Presentation Of Peptide Antigen Via Mhc Class Ii | $1.73 \times 10^{-2}$ | *CD74*, *CTSD*, *PYCARD*, *CTSS*, *LGMN*, *FCER1G* |
| Regulation Of Cell-Substrate Adhesion | $1.77 \times 10^{-2}$ | *LGALS1*, *COL1A1*, *MDK*, *POSTN*, *FN1*, *EMILIN1*, *FLNA*, *LRP1* |
| Lymphocyte Mediated Immunity | $1.78 \times 10^{-2}$ | *CD74*, *C1QB*, *C1QA*, *C1QC*, *C3*, *SERPING1*, *CORO1A*, *B2M*, *FCER1G*, *C1S* |
| Response To Nitrogen Compound | $1.84 \times 10^{-2}$ | *APOE*, *SPARC*, *COL6A1*, *CYBA*, *COL1A1*, *VIM*, *ITGB2*, *COL1A2*, *COL3A1*, *TIMP1*, *LGMN*, *FLNA*, *CD68*, *FCER1G*, *LRP1*, *RARRES2*, *ACTB*, *FBN1* |
| Antigen Processing And Presentation Of Peptide Or Polysaccharide Antigen Via Mhc Class Ii | $2.05 \times 10^{-2}$ | *CD74*, *CTSD*, *PYCARD*, *CTSS*, *LGMN*, *FCER1G* |
| Neuron Death | $2.07 \times 10^{-2}$ | *APOE*, *C1QA*, *MDK*, *GPNMB*, *ITGB2*, *TYROBP*, *CORO1A*, *LGMN*, *SERPINF1*, *LRP1* |
| Cell Development | $2.14 \times 10^{-2}$ | *APOE*, *C1QA*, *LGALS1*, *COL6A1*, *SFRP2*, *MDK*, *POSTN*, *VIM*, *C3*, *CTHRC1*, *FN1*, *TYROBP*, *PSAP*, *SEMA3F*, *COL3A1*, *COL6A2*, *FLNA*, *GFRA1*, *B2M*, *COMP*, *SERPINF1*, *INPP5J*, *LRP1*, *ACTB*, *COL6A3*, *FBN1* |
| Humoral Immune Response | $2.17 \times 10^{-2}$ | *C1QB*, *C1QA*, *C1QC*, *A2M*, *LYZ*, *C3*, *FAU*, *SERPING1*, *C1S*, *RARRES2* |
| Regulation Of Cellular Component Organization | $2.26 \times 10^{-2}$ | *APOE*, *APOC1*, *CAPG*, *LGALS1*, *SPARC*, *CYBA*, *SFRP2*, *AEBP1*, *MDK*, *POSTN*, *MUC1*, *VIM*, *ITGB2*, *C3*, *FN1*, *TYROBP*, *PYCARD*, *SEMA3F*, *CORO1A*, *EMILIN1*, *FLNA*, *DCN*, *B2M*, *CD14*, *SERPINF1*, *INPP5J*, *LRP1*, *SPI1* |
| Response To Growth Factor | $2.51 \times 10^{-2}$ | *SPARC*, *SFRP2*, *COL1A1*, *LUM*, *POSTN*, *COL1A2*, *COL3A1*, *CORO1A*, *LGMN*, *EMILIN1*, *DCN*, *COMP*, *VSTM2A*, *FBN1* |
| Regulation Of Proteolysis | $2.55 \times 10^{-2}$ | *APOE*, *SFRP2*, *A2M*, *CTSD*, *C3*, *FN1*, *PYCARD*, *SERPING1*, *TIMP1*, *LGMN*, *SERPINF1*, *LRP1*, *COL6A3*, *PSMB8* |
| Regulation Of Cytokine Production | $2.55 \times 10^{-2}$ | *CD74*, *CYBA*, *LUM*, *MDK*, *GPNMB*, *POSTN*, *C3*, *FN1*, *TYROBP*, *PYCARD*, *B2M*, *CD14*, *FCER1G*, *LRP1* |

| Biological process | Adjusted *p*-value | Intersecting genes |
|---|---|---|
| Immunoglobulin Mediated Immune Response | $2.65 \times 10^{-2}$ | *CD74*, *C1QB*, *C1QA*, *C1QC*, *C3*, *SERPING1*, *FCER1G*, *C1S* |
| Positive Regulation Of Chemokine (C-X-C Motif) Ligand 2 Production | $2.73 \times 10^{-2}$ | *CD74*, *POSTN*, *LRP1* |
| B Cell Mediated Immunity | $2.92 \times 10^{-2}$ | *CD74*, *C1QB*, *C1QA*, *C1QC*, *C3*, *SERPING1*, *FCER1G*, *C1S* |
| Integrin-Mediated Signaling Pathway | $3.02 \times 10^{-2}$ | *ITGB2*, *TYROBP*, *COL3A1*, *TIMP1*, *FLNA*, *FCER1G* |
| Positive Regulation Of Innate Immune Response | $3.08 \times 10^{-2}$ | *CYBA*, *MUC1*, *ITGB2*, *CTSB*, *PYCARD*, *CTSS*, *LGMN*, *CD14*, *FCER1G*, *PSMB8* |
| Innate Immune Response-Activating Signal Transduction | $3.40 \times 10^{-2}$ | *CYBA*, *MUC1*, *ITGB2*, *CTSB*, *CTSS*, *LGMN*, *CD14*, *FCER1G*, *PSMB8* |
| Regulation Of Cell Development | $3.40 \times 10^{-2}$ | *APOE*, *LGALS1*, *SFRP2*, *MDK*, *POSTN*, *VIM*, *FN1*, *TYROBP*, *SEMA3F*, *COL3A1*, *FLNA*, *B2M*, *SERPINF1*, *INPP5J*, *LRP1*, *FBN1* |
| Negative Regulation Of Cellular Component Movement | $3.78 \times 10^{-2}$ | *APOE*, *CD74*, *SFRP2*, *SEMA3F*, *COL3A1*, *TIMP1*, *EMILIN1*, *DCN*, *SERPINF1*, *LRP1* |
| Response To Lipoprotein Particle | $4.03 \times 10^{-2}$ | *APOE*, *ITGB2*, *CD68*, *FCER1G* |
| Regulation Of Angiogenesis | $4.04 \times 10^{-2}$ | *SPARC*, *SFRP2*, *MDK*, *GPNMB*, *ITGB2*, *C3*, *EMILIN1*, *DCN*, *SERPINF1*, *HSPG2* |
| Positive Regulation Of Phagocytosis | $4.18 \times 10^{-2}$ | *CYBA*, *C3*, *PYCARD*, *FCER1G*, *LRP1* |
| Bone Morphogenesis | $4.54 \times 10^{-2}$ | *COL6A1*, *SFRP2*, *COL1A1*, *COL6A2*, *COMP*, *COL6A3* |
| Positive Regulation Of Response To Biotic Stimulus | $4.70 \times 10^{-2}$ | *CYBA*, *MUC1*, *ITGB2*, *CTSB*, *PYCARD*, *CTSS*, *LGMN*, *CD14*, *FCER1G*, *PSMB8* |
| Cellular Response To Amino Acid Stimulus | $4.82 \times 10^{-2}$ | *COL6A1*, *CYBA*, *COL1A1*, *COL1A2*, *COL3A1* |
| Negative Regulation Of Complement Activation, Lectin Pathway | $4.89 \times 10^{-2}$ | *A2M*, *SERPING1* |
| Regulation Of Complement Activation, Lectin Pathway | $4.89 \times 10^{-2}$ | *A2M*, *SERPING1* |
| Negative Regulation Of Locomotion | $4.91 \times 10^{-2}$ | *APOE*, *CD74*, *SFRP2*, *SEMA3F*, *COL3A1*, *TIMP1*, *EMILIN1*, *DCN*, *SERPINF1*, *LRP1* |
